# Modeling 2D Spatio-Tactile Population Receptive Fields of the Fingertip in Human Primary Somatosensory Cortex

**DOI:** 10.1101/2025.05.24.655840

**Authors:** Susanne Stoll, Falk Luesebrink, D. Samuel Schwarzkopf, Hendrik Mattern, Peng Liu, Johanna Noelle, Esther Kuehn

## Abstract

Tactile fingertip sensations are critical for everyday life. Accordingly, tactile fingertip maps have been extensively studied in human primary somatosensory cortex. However, the fine-grained functional architecture of these maps remains largely unknown. To uncover this architecture, we sought to estimate 2D spatio-tactile population receptive fields (pRFs) of the tip of the index finger in human Brodmann area 3b (BA3b). Using functional magnetic resonance imaging at 7T and submillimeter resolution along with prospective motion correction, we recorded brain responses whilst participants sensed a row of vibrotactile pins sweeping along cardinal axes over a portion of the fingertip. To estimate pRF position and size, we initially fit a 2D Gaussian pRF model to the data, which, however, produced largely implausible pRF estimates. Simulations indicated that this likely occurred because the size of pRFs in BA3b surpasses the portion of the fingertip we stimulated, resulting in an incomplete mapping of pRFs. To address this issue, we constrained the fitting procedure and refined the 2D Gaussian pRF model by keeping pRF size constant. Our results for pRF position then revealed that the ulnar-to-radial axis spanning the fingertip maps onto a superior-to-inferior axis in BA3b. Both the putatively large pRF size (relative to the mapping area) and the pRF position gradient we uncover here appear compatible with receptive field properties quantified in monkeys. Our study provides the first comprehensive investigation into the fine-grained functional architecture of human fingertip maps and brings us one step closer to a thorough understanding thereof.

## 1 Introduction

Day in, day out, we use our fingertips for a variety of touch-based tasks, such as twisting a cap, pressing a lever, tapping on a pad, popping a pimple, or stroking someone else’s face. In fact, by guiding both our perception and motor actions, tactile fingertip sensations render our interaction with the world incredibly effective. This becomes most evident when such sensations are suddenly impaired following (temporary) paresthesia, disease, injury, or amputation (e.g., Gorniak et al., 2014; Kawaiah et al., 2020; Nora et al., 2004). Yet, despite the functional significance of fingertip sensations for our daily lives, we know surprisingly little about how tactile input to the fingertips is represented in the human brain. More precisely, although we know that human primary somatosensory cortex (SI) represents our fingertips in a somatotopic fashion (for a review, see Janko et al., 2022), we lack insights into the fine-grained architecture of an individual fingertip representation.

The building blocks of this fine-grained architecture are tactile neurons and their 2D spatial receptive fields – the region on the skin a tactile neuron is sensitive to. To investigate spatial receptive field properties of the fingertips at the level of neuronal populations, a plethora of studies has combined functional magnetic resonance imaging (fMRI) at 7T with phase-encoding analyses (Engel, 2012; Engel et al., 1994; Kuehn et al., 2018; P. Liu et al., 2021; Puckett & Sanchez-Panchuelo, 2023; Puckett et al., 2020; Sanchez-Panchuelo et al., 2010; Sanchez-Panchuelo et al., 2018). This is because fMRI at an ultra-high field strength (≥ 7T) allows for high image resolution whilst ensuring a good signal-to-noise ratio of the fMRI response, which is thought to help resolve small somatotopic gradients in individual brains (Puckett & Sanchez-Panchuelo, 2023).

For phase-encoded fingertip mapping, participants’ fingertips (or a subregion thereof) are typically successively stimulated vibro-tactilely while voxel-wise blood-oxygen-level-dependent (BOLD) time series are recorded. This is also referred to as between-finger(tip) mapping. Assuming neurons in different voxels have receptive fields preferentially encoding different fingertips, the stimulus-evoked responses in these time series should vary systematically in their delay. As such, a sine wave can be fit to each time series to obtain a phase estimate reflecting this delay^1^ and thus which voxel responds preferentially to which fingertip (Engel, 2012; Engel et al., 1994; Puckett & Sanchez-Panchuelo, 2023). Using this approach, the existence of between-fingertip maps has been repeatedly demonstrated, where the tips of the thumb, index, middle, ring, and little finger are typically represented somatotopically along an inferior-to-superior axis in Brodmann area (BA) 3b, 1, and 2 in SI (Kuehn et al., 2018; P. Liu et al., 2021; Puckett et al., 2020; Sanchez-Panchuelo et al., 2010; Sanchez-Panchuelo et al., 2018). Notably, such maps have also been reported for other experimental designs, analysis techniques, stimulus types, and lower field strengths (3T, 4T) as well as for multi-focal finger stimulation and finger tapping (for a review, see Janko et al., 2022).

A more explicit way to investigate spatial receptive field properties at the level of neuronal populations is fMRI-based population receptive field (pRF) modeling (Dumoulin & Knapen, 2018; Dumoulin & Wandell, 2008). A pRF refers to the aggregate receptive field properties of a population of neurons within a voxel (see Figure 1; Dumoulin & Knapen, 2018; Dumoulin & Wandell, 2008). Similar to phase-encoding analyses (Engel, 2012; Engel et al., 1994), pRF modeling has been pioneered in the field of vision science to study retinotopic maps in the human brain (Dumoulin & Knapen, 2018; Dumoulin & Wandell, 2008). However, unlike phase-encoding analyses, pRF modeling typically uses biological encoding models inspired by invasive electrophysiology to estimate pRF properties (Dumoulin & Knapen, 2018; Naselaris et al., 2011). As such, pRF modeling makes explicit assumptions about how a stimulus interacts with the pRF of a voxel over time to obtain predicted BOLD time series, which are then compared to actually observed BOLD time series (Dumoulin & Knapen, 2018; Dumoulin & Wandell, 2008). A perfect pRF model can thus – at least in theory – provide an exhaustive functional characterization of a certain brain region (Naselaris et al., 2011).

**Figure 1.**
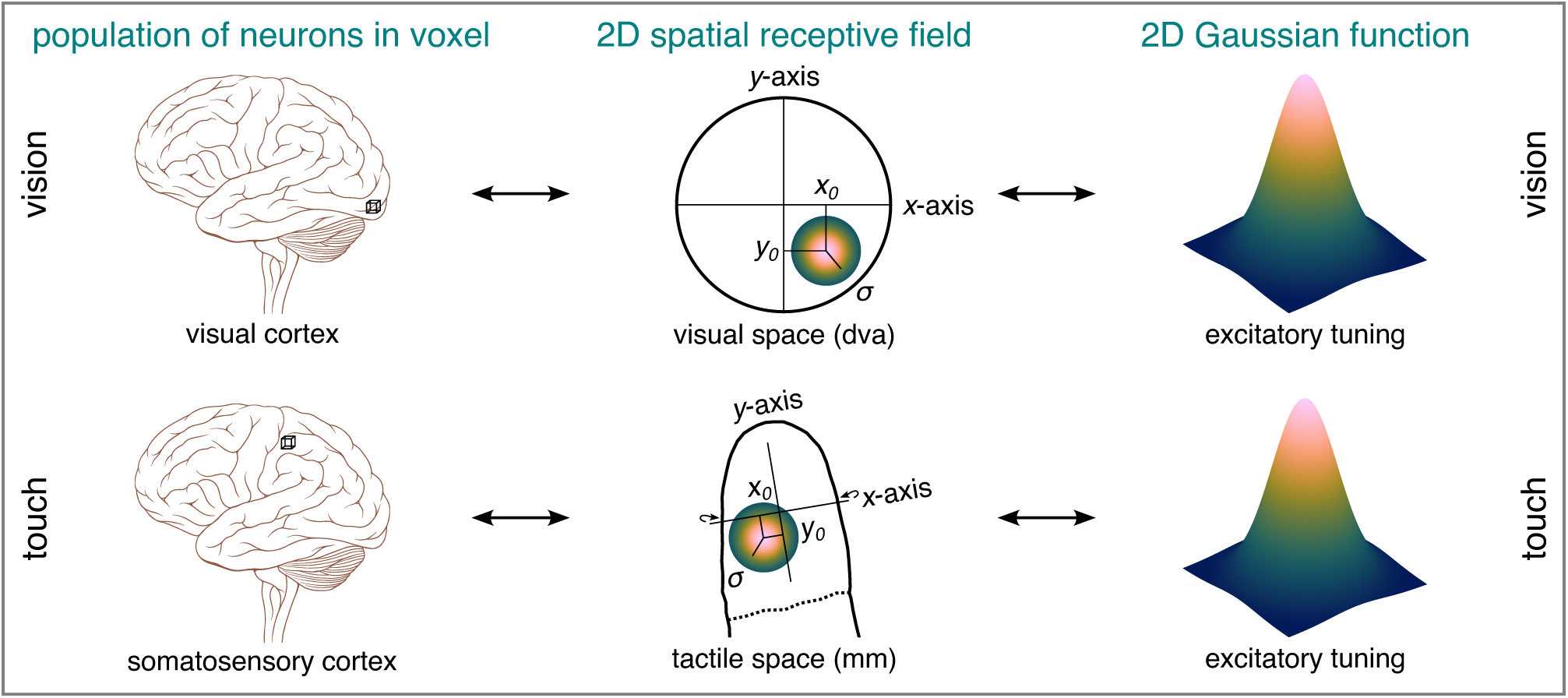
2D spatial pRF. A 2D spatio-visual pRF refers to the 2D region on the retina and thus visual space a population of neurons within a voxel (and most likely beyond) is sensitive to when visually stimulated. A 2D spatio-tactile pRF of the fingertip refers to the 2D region on the fingertip skin and thus tactile space a population of neurons within a voxel (and most likely beyond) is sensitive to when tactilely stimulated. Both types of pRFs can be modeled as an 2D excitatory Gaussian function, where *x*_0_ and *y*_0_ coordinates represent its center position and *σ* its size in sensory space (i.e., a Cartesian coordinate system). Wrap-around arrows indicate an *x*-axis that is slightly wrapped around the fingertip. pRF = population receptive field. dva = degrees of visual angle. The brain insets are adapted from *Brain human lateral view*, by P. J. Lynch, 2006, Wikimedia (https://commons.wikimedia.org/wiki/File:Brain_human_lateral_view.svg). CC BY 2.5.

In vision science, pRFs are typically modeled as a 2D excitatory Gaussian function, which allows the center position (*x*_0_, *y*_0_) and size (*σ*) of a pRF to be estimated in 2D sensory space (see Figure 1; Dumoulin & Knapen, 2018; Dumoulin & Wandell, 2008). To adapt 2D pRF modeling to the case of 1D between-fingertip stimulation as described above, the 2D Gaussian pRF model can be either constrained by keeping one dimension constant (e.g., fixing *y*_0_) or reduced to a 1D Gaussian pRF model (e.g., removing *y*_0_). A series of high-resolution 7T-fMRI studies adopting such approaches did not only show that pRF position maps in SI are reminiscent of between-fingertip maps typically obtained via phase estimates, but also that overall pRF size might differ between different digits, although individual results did not always converge (P. Liu et al., 2021; Puckett et al., 2020; Schellekens et al., 2021). Similar results have been obtained for finger movements (Schellekens et al., 2018), multi-focal finger stimulation (Asghar et al., 2023), more complex types of adapted 2D pRF modeling (Puckett et al., 2020), and various types of full 2D pRF modeling based on stimulation between and within fingers (Asghar et al., 2023; Wang et al., 2021). What this line of research leaves completely unaddressed, though, is how neighboring points on a given fingertip and thus the skin’s 2D receptor sheet maps onto human SI (see Figure 1; Sereno et al., 2022).

Unlike research in humans, research in macaques and/or owl monkeys involving invasive singleneuron or multi-unit recordings started tackling this question a while ago (Merzenich et al., 1978; Nelson et al., 1980; Sur et al., 1980). Using dense tactile stimulation delivered via fine glass probes, this line of work revealed that 2D excitatory receptive fields are organized so that the ulnar-to-radial and the distal-to-proximal axis spanning a monkey’s finger(tip) are represented along a superior-to-inferior and anterior-to-posterior axis in BA3b, respectively (Merzenich et al., 1978; Nelson et al., 1980). It furthermore indicated that the area of excitatory receptive fields in BA3b can range from a subportion of to almost the entire fingertip (Merzenich et al., 1978; Nelson et al., 1980; Sur et al., 1980).

Interestingly, a series of human fMRI studies differing along various methodological dimensions (e.g., experimental design, analysis technique, stimulus type, field strength, image resolution, digits of interest) has already investigated the representation of the distal-to-proximal axis along the fingers in BA3b. This is also referred to as within-finger mapping. Whereas some findings point to an orderly mapping just like in monkeys (Blankenburg et al., 2003; Choi et al., 2016; Sanchez-Panchuelo et al., 2012; Śanchez-Panchuelo et al., 2014), others seem less clear (Asghar et al., 2023; P. Liu et al., 2025; Overduin & Servos, 2004, 2008; Schweisfurth et al., 2014, 2015; Wang et al., 2021). In these studies, tactile stimulation progressed successively within individual or multiple fingers, sometimes combined with stimulation between fingers, and the data were analyzed using conventional general linear models (GLMs), phase-encoding analyses, 1D pRF modeling, adapted 2D pRF modeling, and various types of full 2D pRF modeling. However, the granularity of stimulation in these studies was coarse (no more than approximately two effective locations per phalange) and the ulnar-to-radial axis within individual fingers was not mapped, rendering it impossible to fully resolve the fine-grained architecture of an individual fingertip representation.

In the present exploratory study, we therefore sought to estimate fine-grained, 2D spatio-tactile pRFs of the fingertip in human BA3b (see Figure 1). To maximize sensitivity, we combined 7T-fMRI at submillimeter resolution with prospective motion correction (PMC; Maclaren et al., 2012; Stucht et al., 2015) and precision neuroscience (Gordon et al., 2017). Whereas PMC refers to correcting head motion at the time of data acquisition, precision neuroscience refers to collecting a large amount of data per individual for a detailed characterization of individual brains. To map 2D spatio-tactile pRFs, we recorded BOLD responses whilst participants sensed a fine-grained row of vibrotactile pins moving along cardinal axes over a portion of the tip of the index finger. To estimate 2D spatio-tactile pRFs, we fit an 2D excitatory Gaussian pRF model to the data. To refine our modeling procedure and investigate the reliability, explanatory power, computational validity (accuracy), and meaningfulness of our results, we furthermore performed reliability, cross-validation, and simulation analyses. Our study provides the first comprehensive investigation into the intricate architecture of an individual fingertip representation in the human brain.

## 2 Methods

### 2.1 Participants

The study sample comprised 3 healthy participants (2 female, 1 male; age range: 27-39 years; all right-handed; all authors) referred to as subject 01, 02, and 03 (Sub-01, Sub-02, and Sub-03). Two of them were highly experienced magnetic resonance imaging (MRI) participants. None of them were näıve as to the purpose of the experiment. All participants were screened for contraindications for 7T-(f)MRI and PMC, gave written informed consent prior to participating in the experiment, and reported intact sensory and motor function of their hands. All participants furthermore gave explicit written informed consent that their pseudonymized brain data can be shared publicly without any form of defacing. Experimental procedures were approved by the Ethics Committee of the Ottovon-Guericke University (OVGU) Magdeburg.

### 2.2 Apparatus

A whole-body MAGNETOM 7T-MRI scanner (7T Classic; Siemens Healthineers, Erlangen, Germany) along with a 32-channel head coil (Nova medical, Wilmington, MA, USA) was used to acquire functional and anatomical images at the MRI Research Infrastructure of the OVGU Magdeburg. As a visual anchor (in an otherwise dark and tiny space), a blank display was projected onto a screen at the back of the MRI scanner. Participants viewed the screen through a mirror mounted onto the head coil. Participants’ head was stabilized via foam padding wrapped around the entire back part of their head (from cheek bone to cheek bone). An MRI-compatible piezostimulator developed by QuaeroSys (https://www.quaerosys.com/) was used to stimulate a portion of the tip of the left index finger (glabrous skin) via a single stimulation module comprising 16 vibrotactile pins. These pins had a pointy pinhead (maximal diameter: ∼1 mm; distance from pin center to pin center: ∼2.5 mm), were arranged as a 4×4 array (see Figure 2), and passed through a surface containing 16 circular apertures. The piezostimulator had a voltage output of 4096 levels representing the possible range of pin heights. A dark surgical marker was used to draw the 4×4 pin array onto participants’ fingertips (see 2.4 Procedure) and an elongated cushion to stabilize the left hand.

**Figure 2.**
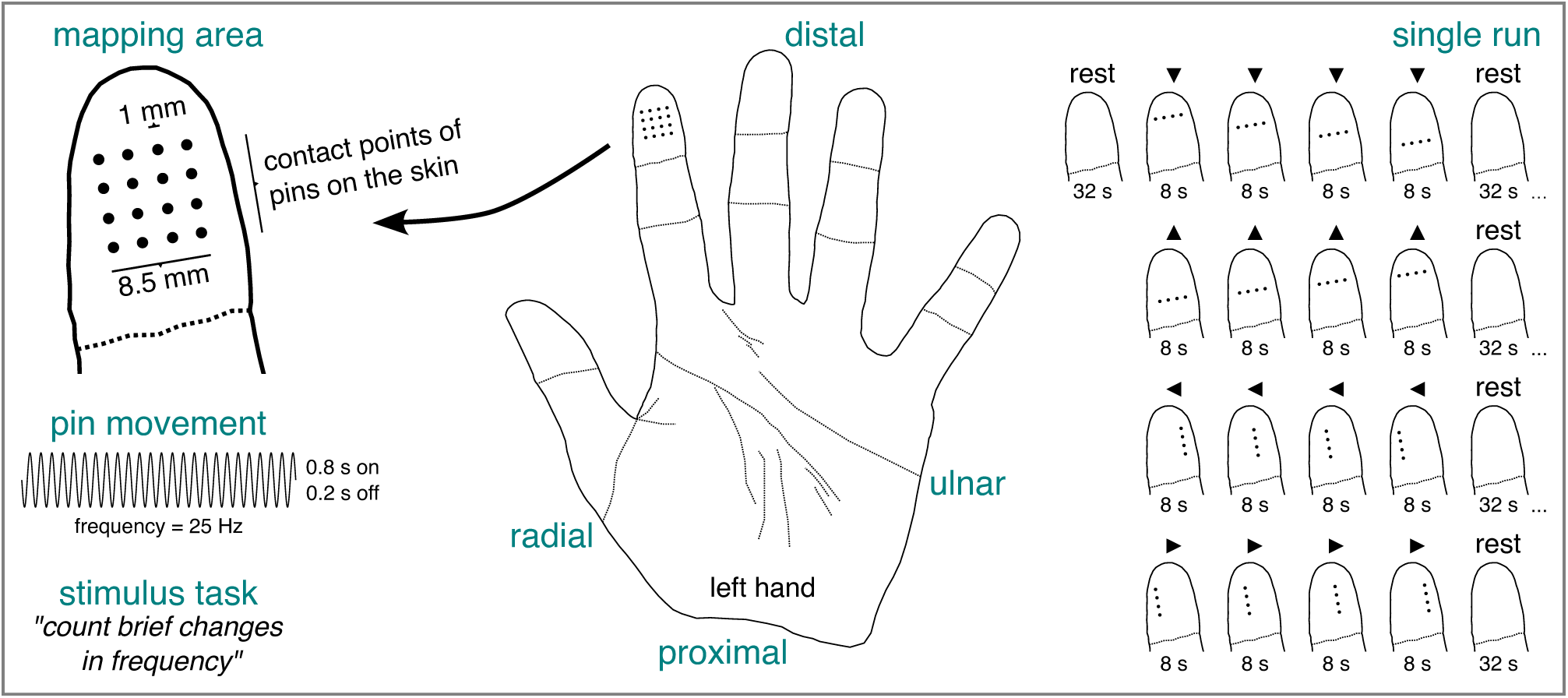
Spatio-tactile pRF mapping experiment. The mapping area consisted of an array of vibrotactile pins placed on a portion of the tip of the left index finger. In any given run, participants sensed a row of pins moving along cardinal axes interspersed with baseline (rest) periods. The pins oscillated according to a sine wave with alternating on and off periods. Participants were required to count brief changes in frequency of the sine wave throughout a given run. Black arrowheads indicate the movement direction of the pins. pRF = population receptive field.

Head motion was corrected online via a PMC system consisting of an MRI-compatible camera (Metria Innovation, Milwaukee, Wisconsin, USA) with a lighting unit illuminating a marker with Moiŕe patterns (size: 15×15 mm^2^). The camera including lighting unit was mounted onto the MRI bore’s ceiling above the head coil and tracked the marker with a sampling rate of 86 Hz. The marker was attached to the extending bit of an MRI-compatible customized mouthpiece that was worn by participants during scanning. The customized mouthpieces were manufactured at the University Clinic for Oral and Maxillofacial Surgery in Magdeburg, covered 6 teeth of the upper jaw (i.e., the central and lateral incisors and the canines), and ensured rigid coupling between the marker and participants’ head. The video stream captured by the camera was sent to a dedicated computer that processed the image frames and estimated head motion along 6 degrees of freedom: 3 rotation axes (pitch, roll, yaw) and 3 translation axes (forward-back, up-down, left-right). These estimates were then forwarded to the MRI scanner and converted from the camera’s coordinate system to the scanner’s coordinate system via a precalibrated transformation matrix. The PMC system tracks motion with a precision of ∼ 0.01 mm (translation) and ∼ 0.01^°^ (rotation), which is more than an order of magnitude smaller than the voxel edge length. More details on the PMC system and its validation can be found in Maclaren et al. (2012) and Stucht et al. (2015).

### 2.3 Stimuli

The pRF mapping area consisted of an array of vibrotactile pins that was divided into 4 horizontal and 4 vertical bars, each comprising 4 pins, resulting in 8 stimulus positions. A single bar therefore functioned as the pRF mapping stimulus. The pins of this bar stimulus typically oscillated according to a sine wave with a constant frequency (*A* = 3095; *f* = 25 Hz; *ϕ* = 4.71 rad; for possible pin heights, see 2.2 Apparatus). However, as part of an oddball task (see 2.4 Procedure), the bar stimulus occasionally oscillated according to a sine wave with a lower frequency (*f* = 10 Hz). For an illustration of the pRF mapping area and stimulus, see Figure 2. To account for an accumulation of small temporal delays in the order of several seconds over the course of a full run of the pRF mapping experiment, we occasionally dropped points sampled from the sine wave. The bar stimulus used here can be seen as the tactile equivalent of bar stimuli comprising flickering high-contrast images frequently adopted for visual pRF modeling (e.g., Alvarez et al., 2015; Dumoulin & Wandell, 2008).

### 2.4 Procedure

Adhering to the principles of precision neuroscience (Gordon et al., 2017), we collected a large amount of data for each participant. In particular, the pRF mapping experiment consisted of 4 sessions *à* 10 runs per participant. For each participant, these 4 sessions were split across 2 days (either consecutive or 1 day apart), so that session 1-2 were conducted on the first day and session 3-4 on the second day. There was a break of ∼15-30 min between sessions on the same day, during which participants were taken out of the MRI scanner. At the beginning of each session, a series of prescans (localizer, transmit adjustment, and shimming) was acquired, followed by anatomical data, followed by functional BOLD data for the pRF mapping experiment and a series of additional scans (see 2.5 MRI acquisition).

At the beginning of each pRF mapping run, 4 dummy volumes were acquired. Next, the bar stimulus translated step-by-step along a portion of the tip of the left index finger. Specifically, it translated from distal-to-proximal (32 s), proximal-to-distal (32 s), ulnar-to-radial (32 s), and radial-to-ulnar (32 s) with a baseline interval (32 s) before/after every sweep. During the dummy volumes and the baseline interval, no stimulation was applied. Note that baseline intervals are necessary for estimating large pRFs (Dumoulin & Knapen, 2018; Dumoulin & Wandell, 2008). Each bar position was stimulated for 8 s. To prevent adaptation, pin oscillation was ceased for 0.2 s every 0.8 s. Moreover, to ensure sustained attention, participants were asked to detect 0-10 oddball events throughout each run. They were instructed to count these events and verbally report whether there were more than 5 oddballs at the end of a given run. The number and timing of oddball events in any given run was determined randomly. During an oddball event, the frequency of the sine wave changed briefly for 0.2 s (see 2.3 Stimuli). For an illustration of the pRF mapping experiment, see Figure 2.

Prior to the start of the first scanning session on a given day, the camera had to warm up for ∼1 hr. During this time, we performed 1-2 mock pRF runs. Moreover, before the start of each pRF mapping run, a subset of pins was supposed to rise and fall once and participants were asked to report if this was not the case. These check-ups were conducted to ensure that all components of the stimulation equipment were intact and the experiment could run through properly. Nonetheless, due to the relatively large number of runs collected in any given session, it could happen that the pRF mapping experiment did not run through properly, which was clearly perceivable (e.g., pins stopped oscillating). As such, participants were instructed to report such instances at the end of a given run, in which case this run was repeated. Similarly, a run was repeated whenever participants reported that they were “inattentive” or “sleepy” at the end of the run. Per participant and session, the range of invalid and thus to-be-repeated runs was 0-3.

Prior to the start of session 1, participants performed a training session outside the MRI scanner, where they familiarized themselves with the pRF mapping experiment and the oddball task. All participants were able to correctly report the movement direction of the bar stimulus, to detect the oddball events, and perceive each bar position clearly and distinctly. To ensure consistent placement of the pins in each session, we aligned the lower edge of the surface element of a second (identical) stimulation module with the terminal finger joint, and used a dark surgical marker to draw the 4×4 pin array onto each participant’s fingertip. This happened prior to the start of session 1. If not actively washed off, the drawing could remain on the fingertip for several days, which was the case for each participant. We also took a picture of the drawing prior to session 1 and prior to session 3. Moreover, to ensure that the pins remained in the same position throughout a whole scanning session, the stimulation module was attached to the index finger using medical tape. Participants were furthermore instructed to keep still during scanning and to refrain from moving their hand and fingers in particular. An overview of system environments and general-purpose software tools adopted in the pRF mapping experiment can be found in Table S1. This table also covers visualizations presented in 1 Introduction and 2 Methods.

### 2.5 MRI acquisition

After localizing participants’ brains, we calibrated the transmit voltage, followed by head shimming to make the main magnetic field (B0) more homogenous. Subsequent to these prescans, we acquired whole-brain anatomical images using a T1-weighted 3D magnetization-prepared rapid acquisition with gradient echo (MPRAGE) sequence (Mugler & Brookeman, 1990, see Table 1). We then mapped geometric distortions induced by residual B0 field inhomogeneities via the point-spread function (PSF) method (In & Speck, 2012; In et al., 2015; Zaitsev et al., 2004). Afterwards, we acquired several runs of functional BOLD images for pRF mapping (see 2.4 Procedure). In both cases (geometric distortions and pRF), we used a T2∗-weighted 2D gradient-echo (GE) echo planar imaging (EPI) sequence (see Table 1). Between the first and second run of functional data acquisition, we additionally mapped geometric distortions using the T2∗-weighted 2D GE EPI sequence for acquiring functional data with reversed phase-encoding as well as a T2∗-weighted 2D GE sequence with two different echo times. However, these additional images were only acquired for potential future investigations and not used in the scope of the present study.

**Table 1.** Parameters of Main Scanning Sequences.

| Characteristics | 3D MPRAGE | 2D GE EPI | 2D GE EPI (PSF) |
| --- | --- | --- | --- |
| Number of slices | 224 | 30 | 30 |
| Slice orientation | sagittal | t tilted twds. c | t tilted twds. c |
| Slice order |  | interleaved | interleaved |
| Voxel size | 0.8 mm iso | 0.9 mm iso | 0.9 mm iso |
| TR | 2.5 s | 2 s | 2 s |
| TE | 2.45 ms | 20 ms | 20 ms |
| TI | 1050 ms |  |  |
| Flip angle | 5° | 76° | 90° |
| PI | GRAPPA | GRAPPA | GRAPPA |
| Acceleration factor | 2 | 4 | 4 |
| Ref. lines for PI | 32 | 64 | 64 |
| BW | 320 Hz/pixel | 1072 Hz/pixel | 1072 Hz/pixel |
| FoV | 218×218×179.2 mm | 200×200×27 mm | 200×200×27 mm |
| Echo spacing | 5.8 ms | 1.08 ms | 1.06 ms |
| Image interpolation |  | zero filling |  |
| Phase partial Fourier |  | 5/8 | 5/8 |
| Number of volumes | 1 | 144 | 1 |
| PSF reduced FoV factor |  |  | 4 |
| EPI-PSF acceleration factor |  |  | 1 |
| EPI-PSF reference lines |  |  | 8 |

*Note.* Blank entries denote "not applicable". TR = repetition time. TE = echo time. TI = inversion time. Ref. = reference. PI = parallel imaging. BW = bandwidth. FoV = field-of-view. PSF = point-spread function. GE = gradient-echo. EPI = echo planar imaging. MPRAGE = magnetization-prepared rapid acquisition with gradient echo. iso = isotropic. GRAPPA = generalized autocalibration partial parallel acquisition. t = transversal. twds. = towards. c = coronal.

For acquiring functional images for pRF mapping and mapping geometric distortions, the image slab was centered at the base of the hand knob (i.e., a posteriorly-oriented hook when viewed sagittally; Yousry et al., 1997) and then titled so that the anterior-posterior midline of the slab (the longer line of symmetry) was aligned with an oblique axis at approximately 160/340^°^. Moreover, any acquired dummy volumes (see 2.4 Procedure) were directly discarded by the MRI scanner. As for the MPRAGE sequence, the motion estimates obtained via the PMC system were used to update the field-of-view prior to the excitation of each individual k-space line. As for the EPI sequences, this was done prior to the excitation of all k-space lines pertaining to an individual slice. As such, for both anatomical and functional data, PMC accounted for head motion within but not between runs or sessions. The distortion correction of the functional images using the PSF method was performed online via a fully-automated procedure. All acquired images were exported in DICOM format. The entire sequence protocol is publicly available (see Data and code availability).

### 2.6 Preprocessing

We designed a customized preprocessing pipeline that aligns and coregisters anatomical and functional images across all 4 sessions and constructs an anatomical surface model for each participant without manual intervention. The input data for this pipeline were the anatomical and distortion-corrected functional DICOM images that had been prospectively corrected for within-run motion (see 2.5 MRI acquisition). As a preparatory step, we removed invalid functional runs (see 2.4 Procedure) as well as the first volume of each functional run which was the reference volume for distortion correction. We then converted all remaining functional and anatomical DICOM images to NIfTI format and formatted them according to the Brain Imaging Data Structure (BIDS; Gorgolewski et al., 2016).

As for the BIDS-formatted anatomical images of each session, we subsequently downsampled these to match the resolution of the functional images (interpolation method: cubic), B1-bias-field-corrected them via unified segmentation adjusted for high-resolution 7T data (Ashburner & Friston, 2005; Lüsebrink et al., 2017, 2021), and finally masked them using a deep learning strategy for skullstripping (Hoopes et al., 2022, for more details, see Explanation S1). To obtain an initial masked anatomical template across sessions, we then aligned these masked anatomical images (interpolation method: nearest neighbor; for more details, see Explanation S2) and calculated the median across them (Lüsebrink et al., 2017, 2021). To obtain an unmasked anatomical template across sessions, we furthermore applied the resulting transformation matrices to the unmasked bias-field-corrected anatomical images, thus bringing them in alignment (interpolation method: nearest neighbor), and calculated the median across them (for more details, see Explanation S3). To obtain a final masked anatomical template across sessions (for more details, see Explanation S4), we masked this unmasked anatomical template by (yet again) using a deep learning strategy for skull-stripping (Hoopes et al., 2022). The final masked anatomical template was used for coarse coregistration purposes (see below). The unmasked anatomical template was used for reconstructing an anatomical cortical surface model (Dale et al., 1999; Fischl, 2012; Fischl et al., 1999, for more details, see Explanation S5). As part of the reconstruction process, we adopted (yet again) a deep learning strategy for generating a brain mask (Hoopes et al., 2022). A subset of the reconstructed anatomical surfaces was later converted from surface to MAT format (along with the vertex-wise functional time series, see below). Lastly, we derived cytoarchitectonic labels using a probabilistic atlas (Fischl et al., 2008) including labels for right and left BA3b as well as anatomical labels using the gyral-based Desikan-Killiany cortical atlas including labels for right and left postcentral gyrus, that is, the location of SI (Desikan et al., 2006).

As for the BIDS-formatted functional images of each session, we first averaged across all images of a given run. Next, we generated a brain mask by thresholding these run-wise functional averages, obtaining the largest component (to remove non-brain tissue), and morphologically closing holes (due to noise/imperfect thresholding). Subsequently, we used these run-wise brain masks to mask the run-wise average functional images. All masked run-wise average functional images were then used to generate a masked functional template across sessions by aligning these images (interpolation method: nearest neighbor) and calculating the median across them. The final masked anatomical template (see above) was then coregistered to this masked functional template (interpolation method: nearest neighbor; for more details, see Explanation S6). Next, we applied both resulting types of transformation matrices to every BIDS-formatted functional image of each run in one go (interpolation method: nearest neighbor; for more details, see Explanation S7), therefore aligning them and coregistering them to the final masked anatomical template. More specifically, we used the forward transformation matrices for generating the masked functional template and the inverse of the transformation matrix for coregistering the final masked anatomical template to the masked functional template. This first coarse coregistration step was used as a seed for a second fine coregistration step, where we coregistered every coarsely-coregistered functional image of each run to the unmasked anatomical template (interpolation method: nearest neighbor) using the previously generated cortical surface model and boundary-based registration (BBR; Greve & Fischl, 2009; Polimeni et al., 2018, for more details, see Explanation S8). Lastly, we used the transformation matrices resulting from BBR to project the coarsely-coregistered functional data onto the cortical surface model, thus resampling them to surface vertices. To this end, we used the voxel in the functional image that was located midway (projection fraction = 0.5) between corresponding vertices on the gray-white matter boundary and the pial boundary (for more details, see Explanation S9). Moreover, we modestly smoothed the functional data along the cortical surface (surface FWHM = 1 mm; for more details, see Explanation S10). The resulting vertex-wise functional time series for each run (henceforth fMRI time series) were then converted from MGH to MAT format before undergoing linear detrending and *z*-standardization. Note that fMRI time series can vary substantially between sessions. *Z*-standardization ensures that these time series are comparable and weighted equally when calculating vertex-wise averages (see 2.7.3 PRF modeling).

For an overview of the preprocessing steps, see Figure S1. For an illustration of the outputs of various preprocessing steps, see Video S1-Video S14. For adopted system environments and general-purpose software tools as well as additional notes, see Table S1. The preprocessing pipeline including data after BIDS formatting is publicly available (see Data and code availability).

### 2.7 Data analysis

#### 2.7.1 Functional localization

To localize stimulus-evoked activity in SI, we performed a simple vertex-wise GLM analysis for each participant and brain hemisphere. The input data were the preprocessed vertex-wise fMRI time series for session 1 concatenated across 10 runs. The GLM comprised a constant boxcar regressor for each stimulus position (4 horizontal bars and 4 vertical bars) convolved with a canonical hemodynamic response function (HRF; The FIL Methods Group, 2021) and a constant term for each run. Baseline intervals were modeled implicitly (Pernet, 2014). We computed *t*-statistic maps for the contrast stimulation vs baseline (rest): [1 1 1 1 1 1 1 1 0 0 0 0 0 0 0 0 0 0]. Contrasting stimulation vs rest is known to result in an increase in differential brain activity in stimulated and decrease in non-stimulated cortical sites in contralateral primary sensory areas, such as primary visual cortex (Shmuel et al., 2002; Stoll et al., 2020) and SI (Tal et al., 2017). Ipsilaterally, however, a wide-spread decrease of differential brain activity is typically observed in these areas (Shmuel et al., 2002; Tal et al., 2017).

#### 2.7.2 Delineations

To delineate fingertip clusters, we superimposed the label for postcentral gyrus and BA3b onto the unthresholded *t*-statistic maps rendered onto a spherical cortical surface model of the right brain hemisphere. Next, we visually identified clusters of increased brain activity (*t*-statistic *>* 0) falling entirely into the postcentral gyrus label and falling entirely into or overlapping the BA3b label. We then manually delineated the extent of these clusters.

#### 2.7.3 PRF modeling

To estimate spatio-tactile pRFs per participant and brain hemisphere, we performed a pRF modeling analysis using all 30 runs of session 2-4^2^. Moreover, to conduct a split-half reliability analysis and cross-validation analyses, we repeated this analysis for the 15 odd-numbered and the 15 even-numbered runs of session 2-4. In each case, we first averaged the preprocessed vertex-wise fMRI time series across all runs of interest^3^. We then fit three pRF models to the averaged data. The first model consisted of a 2D excitatory Gaussian function (henceforth 2dg model) with 5 free parameters: pRF center position (*x*_0_, *y*_0_), pRF size (*σ*), pRF baseline (*β*_0_), and pRF amplitude (*β*_1_). The second model consisted of a 2D excitatory Gaussian function where the parameter for pRF size was fixed (*σ* = 8.5 mm; henceforth 2dg-fix model), leaving 4 free parameters (*x*_0_, *y*_0_, *β*_0_, *β*_1_). The third model was a model without any spatial tuning and thus 2 free parameters (*β*_0_, *β*_1_), solely reflecting the presence or absence of tactile stimulation (henceforth onoff model; see also Benson et al., 2018; Prince et al., 2022). For both the 2dg and the 2dg-fix model, we expressed pRF center position and pRF size in physical units (mm) and estimated pRF center position in a Cartesian coordinate system where origin corresponded to the midpoint of the mapping area on the left fingertip (i.e., the pin array). For an illustration of the pRF modeling pipeline, see Figure 3.

**Figure 3.**
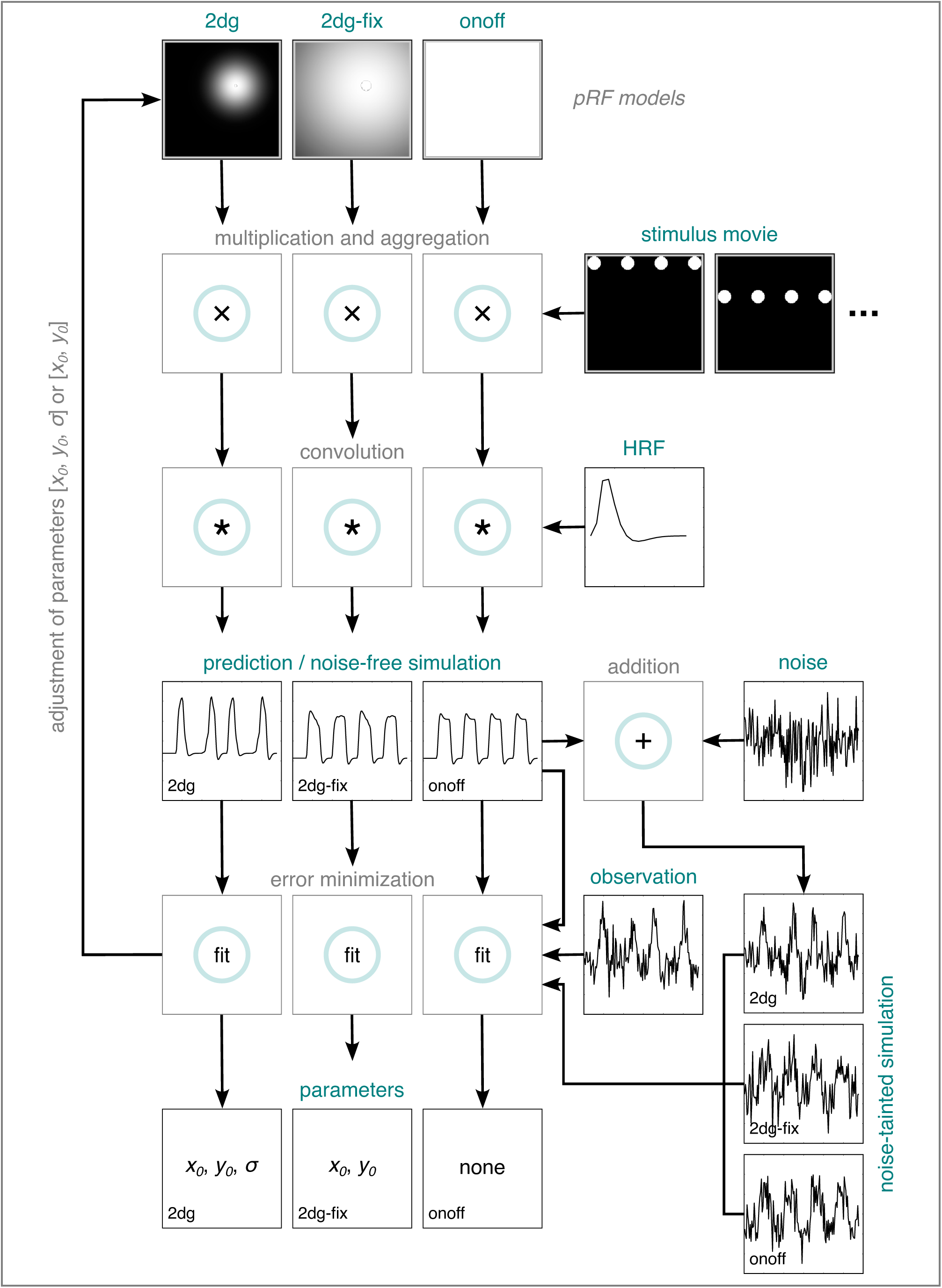
PRF modeling pipeline. *PRF modeling for empirically-observed data*: A pRF model, a stimulus movie, and an HRF are used to predict an fMRI time series, which is then compared to an empirically-observed fMRI time series. This is repeated many times by varying the parameters of the pRF model (if there are any). The parameters producing the best fit (lowest error between predicted and observed time series) are retained. *PRF modeling for simulated data*: The components for predicting an fMRI time series are used to simulate a noise-free fMRI time series. Depending on the purpose of the simulation, this noise-free fMRI time series remains as it is or is repeatedly perturbed by noise. The noise-free or noise-tainted time series are then fed into the pRF modeling pipeline as if they represented empirically-observed data. HRF = hemodynamic response function. 2dg = 2D Gaussian pRF model with unconstrained optimization. 2dg-fix = 2D Gaussian pRF model with parameter for pRF size fixed and constrained grid search. onoff = pRF model without any spatial tuning that simply reflects the presence or absence of tactile stimulation. pRF = population receptive field. fMRI = functional magnetic resonance imaging.

For all three models, we first predicted an fMRI time series. To this end, we calculated an overlap image by multiplying an image of the pRF model and a binarized image of the stimulus per volume (a stimulus movie). These images consisted of a 170×170 pixel matrix, where 1 pixel corresponded to 0.05 mm on the skin. For each overlap image, we then aggregated the pixel values by calculating the mean percent overlap with the image of the pRF model, resulting in a predicted neuronal time series. This neuronal time series was then convolved with a canonical HRF (The FIL Methods Group, 2021, see also 2.7.1 Functional localization), resulting in a predicted fMRI time series. For the 2dg model, we generated 8750 predictions by sampling all putative parameter combinations from a 3D search space for pRF position and size (*x*_0_, *y*_0_ = from −10.625 to +10.625 mm with 25 evenly-spaced values; *σ* = from 0 to 12.75 mm with 15 evenly-spaced values and 0 being excluded). For the 2dg-fix model, we generated 625 predictions by sampling from the same search space but dropping the dimension for pRF size because pRF size was fixed for this model. For the onoff model, we generated a single prediction. We then determined the Pearson correlation between the predicted and observed fMRI time series. For the 2dg and the 2dg-fix model, we subsequently selected the combination of parameter values showing the largest goodness-of-fit (squared Pearson correlation, *R*^2^) as final pRF estimates. Only for the 2dg model, this constrained grid search fit was followed by an unconstrained optimization procedure taking the grid search fits as seeds and using a Nelder-Mead algorithm to further maximize the goodness-of-fit (Lagarias et al., 1998; Nelder & Mead, 1965). This is because the 2dg model with unconstrained optimization resulted in largely implausible parameter estimates (see 3.2 Fingertip pRFs in BA3b are larger than the mapping area). To address this issue, we introduced the 2dg-fix model with a constrained grid search and based all further analyses on this model. And finally, to estimate *β*_1_ and *β*_0_, we performed linear regression between the observed and the predicted fMRI time series representing the final fit for each model. Given that we linearly detrended and *z*-standardized the former and due to the way we calculated the latter, the *β*_1_ and *β*_0_ estimates are not directly interpretable and simply reflect offset and scaling factors, respectively, used to adjust the fitted fMRI time series.

To validate the explanatory power of our results, we conducted a 2-fold-cross-validation analysis where the goodness-of-fit for pRF parameter estimates obtained for the odd runs was calculated using the even runs (first cross-validation fold) and vice versa (second cross-validation fold) for both the 2dg-fix model and the onoff model. To investigate whether the 2dg-fix model has superior explanatory power over the onoff model, we compared the cross-validated goodness-of-fit (*cR*^2^) per vertex for this model to the cross-validated goodness-of-fit for the onoff model. This is possible because the cross-validation procedure we apply here automatically accounts for model complexity. And lastly, to determine how much the overall explanatory power varies between participants and models, we extracted all vertices falling within a given delineated BA3b cluster (see 2.7.2 Delineations) and determined the median cross-validated goodness-of-fit for each participant and model.

#### 2.7.4 Simulations

To investigate the implausibility, meaningfulness, and computational validity (e.g., Lerma-Usabiaga et al., 2020; Senden et al., 2014; Zeidman et al., 2018) of our pRF estimates, we performed a series of simulations. The basic principle underlying these simulations consists in simulating a noise-free fMRI time series for a specific pRF model by following the steps for predicting an fMRI time series (see 2.7.3 PRF modeling and Figure 3). Depending on the type of simulation, this time series either remains as it is or is perturbed by random Gaussian noise and *z*-scored afterwards, resulting in a noise-tainted fMRI time series. Next, a specific pRF model is fit to the simulated noise-free or noise-tainted time series by feeding the time series into the pRF modeling pipeline as if it represented truly observed data (see 2.7.3 PRF modeling and Figure 3).

##### Implausibility

To investigate why fitting a 2dg model with unconstrained optimization resulted in implausible pRF estimates (see 3.2 Fingertip pRFs in BA3b are larger than the mapping area), we simulated how well we can recover the parameters of ground-truth pRFs varying in size when we add or do not add noise to the simulated data. This allowed us to test for the hypothesis that for large pRFs surpassing the mapping area, there is not enough unique information left in a noise-tainted time series to keep the parameter estimates within a plausible range. In a noise-free time series, however, this information is present, resulting in accurate recovery irrespective of pRF size. As such, using the 2dg model, we first simulated noise-free time series for ground-truth pRFs with a center position at origin (*x*_0_ = 0 mm; *y*_0_ = 0 mm) and 6 different sizes (*σ* = 1.0625, 2.125, 3.1875, 4.25, 6.375, 8.5 mm). We then perturbed each of these time series 100000 times with random Gaussian noise (*σ_noise_* = 2, expressed in units of mean percent overlap with the pRF model). Next, we fit a 2dg model with unconstrained optimization to both the noise-free and noise-tainted time series. For each ground-truth pRF in the noise-tainted case, we then calculated the median as well as the 1*^st^* and 99*^th^* percentile over the 100000 repeats, individually for each pRF parameter (*x*_0_, *y*_0_, *σ*).

##### Meaningfulness

To investigate whether the observed differences in cross-validated goodness-of-fit between the 2dg-fix and the onoff model are meaningful, we simulated a null distribution. To this end, we used the onoff model to simulate a noise-free time series for a pRF with no spatial tuning. We then perturbed this time series 100000 times with random Gaussian noise (*σ_noise_* = 2, expressed in units of mean percent overlap with the pRF model). We did this twice to generate an odd and an even dataset. Just like for the empirical data (see 2.7.3 PRF modeling), we fit both the 2dg-fix model with constrained grid search and the onoff model to the simulated noise-tainted time series of each set and determined the cross-validated goodness-of-fit for each model via a 2-fold cross-validation analysis. We then calculated the difference in cross-validated goodness-of-fit between the 2dg-fix and the onoff model. Using the difference distribution for each cross-validation fold, we determined *p*-values for a range of small critical difference values (0.01, 0.02, 0.03). To this end, we counted the number of differential values being equal or surpassing our critical value and divided this number by the overall number of differential values. This gives an indication of how probable it is to observe a certain small critical difference if the null hypothesis were true. Finally, just like for the empirical data, we used the simulated odd and even dataset to calculate the median cross-validated goodness-of-fit for both fitted models via a 2-fold cross-validation analysis (see 2.7.3 PRF modeling). This allowed us to check that our simulations approximate empirical estimates of median cross-validated goodness-of-fit, indicating that the level of noise we introduced to perturb noise-free time series was adequate.

The methodological details for simulations testing for the computational validity of the 2dg-fix model with constrained grid search can be found in S1.2.1 Simulations. The entire data analysis pipeline including empirical and simulated data as well as associated visualizations is publicly available (see Data and code availability). An overview of adopted system environments and general-purpose software tools can be found in Table S1.

## 3 Results and interim discussion

### 3.1 Vibrotactile stimulation produces fingertip clusters in BA3b

Figure 4 shows *t*-statistic maps reflecting differential brain activity for the contrast stimulation vs baseline (rest) per participant and brain hemisphere. For each participant, we were able to identify a cluster of increased differential brain activity falling entirely into postcentral gyrus and thus the location of SI and falling entirely into or overlapping right BA3b. These BA3b clusters were small and surrounded by decreases in differential brain activity. Decreases in differential brain activity were also evident for each participant in left BA3b. Overall, this pattern of results is compatible with previous work (Tal et al., 2017) and suggests that we successfully induced responses in right BA3b corresponding to tactile stimulation of the left fingertip – a prerequisite for pRF modeling.

**Figure 4.**
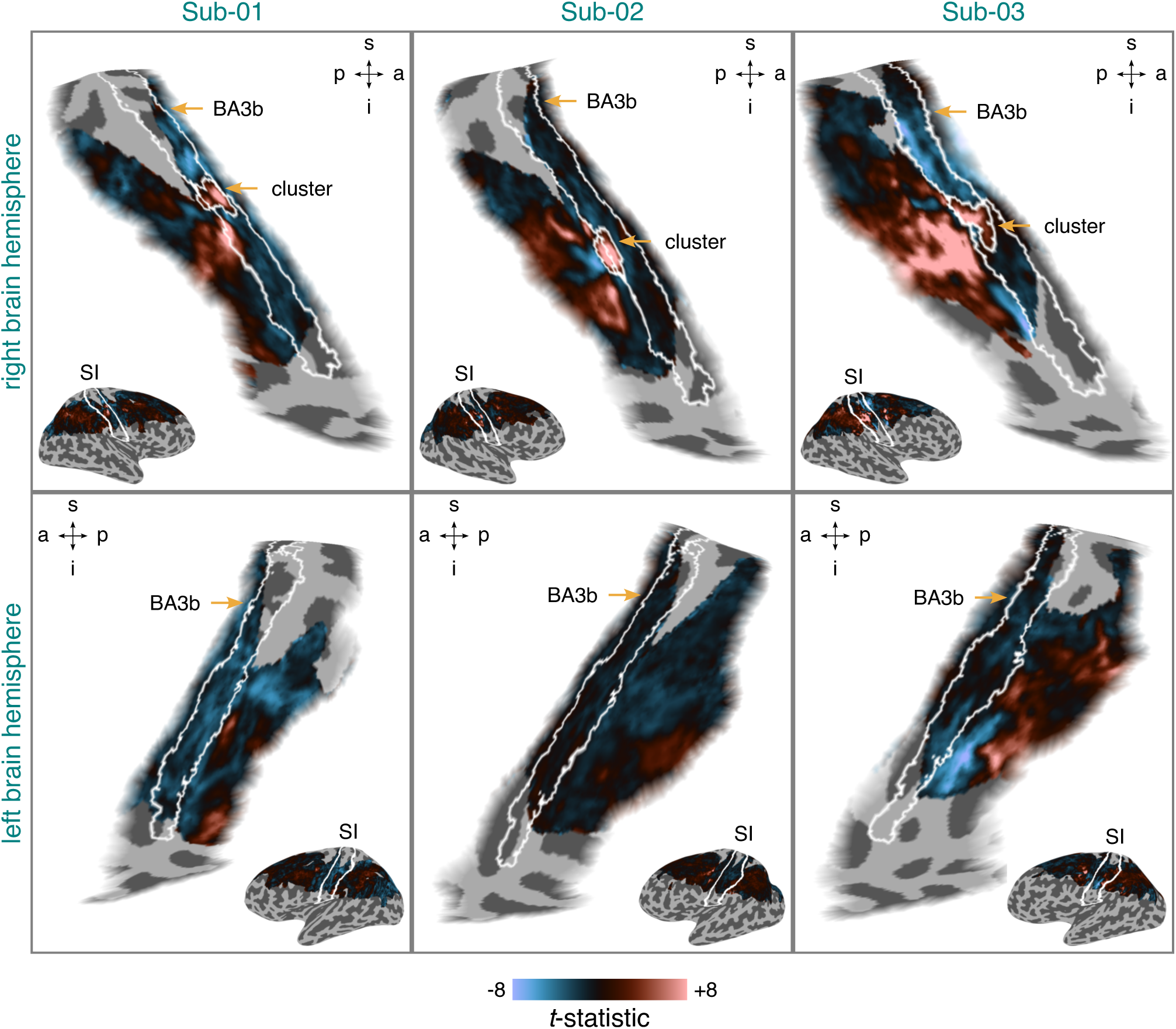
Differential brain activity for stimulation vs baseline (rest) per participant and brain hemisphere. The *t*-statistic maps are based on data from session 1. They were projected onto an inflated cortical surface model where dark gray regions represent sulci and light gray regions gyri. *T*-statistics surpassing a value of ±8 were set to this value. BA3b = Brodmann area 3b. SI = primary somatosensory cortex. s = superior. p = posterior. i = inferior. a = anterior. Sub-01, Sub-02, Sub-03 = subject 01, 02, and 03.

### 3.2 Fingertip pRFs in BA3b are larger than the mapping area

Fitting a 2dg model with unconstrained optimization to all runs resulted in pRF parameter estimates (*x*_0_, *y*_0_, *σ*) in the range of a fraction of a millimeter to several thousand kilometers for the identified BA3b cluster across participants^4^. Extremely large pRF estimates are evidently implausible as no human body could possibly accommodate them. An inspection of the empirically-observed time series suggested that these implausible pRF estimates might be due to broadly-peaked stimulus-evoked responses. Figure 8,Figure S6, and Figure S8 illustrate this for two example vertices based on all runs for each participant. For ordered pRF mapping designs, such as the one adopted here, broad peaks are characteristic of large pRFs, such as pRFs whose size encompasses a large portion of the mapping area or goes well beyond it. Due to the noise being present in empirical time series, broad peaks might be difficult to identify and different broadly-peaked time series difficult to distinguish from one another. This in turn might complicate the unconstrained optimization procedure and result in implausible pRF parameter estimates. To test for this hypothesis, we simulated noise-free and noise-tainted time series for ground-truth pRFs varying in size using a 2dg model and fit a 2dg model with unconstrained optimization to the data.

Figure 5 shows the simulated noise-free time series. This illustration highlights that even without considering the effect of noise, larger pRFs surpassing the mapping area are associated with broader-peaked and less distinctive stimulus-evoked responses. Figure 6 (a.) shows the corresponding recovered pRF parameter estimates (*x*_0_, *y*_0_, *σ*). For all simulated pRF sizes, we were able to closely recover the ground truth. Figure 6 (b.) shows the recovered pRF parameter estimates when noise was added to the simulated data. For pRFs within the mapping area, we were able to closely recover the ground truth using the median (apart from for the smallest pRF). However, as pRF size increased, the upper and lower percentiles became more extreme and thus the range of possible pRF parameter estimates. For pRFs surpassing the mapping area, we were still able to recover the ground truth using the median, albeit less precisely. However, the percentiles of the recovered pRF parameter estimates now indicated an implausibly large range.

**Figure 5.**
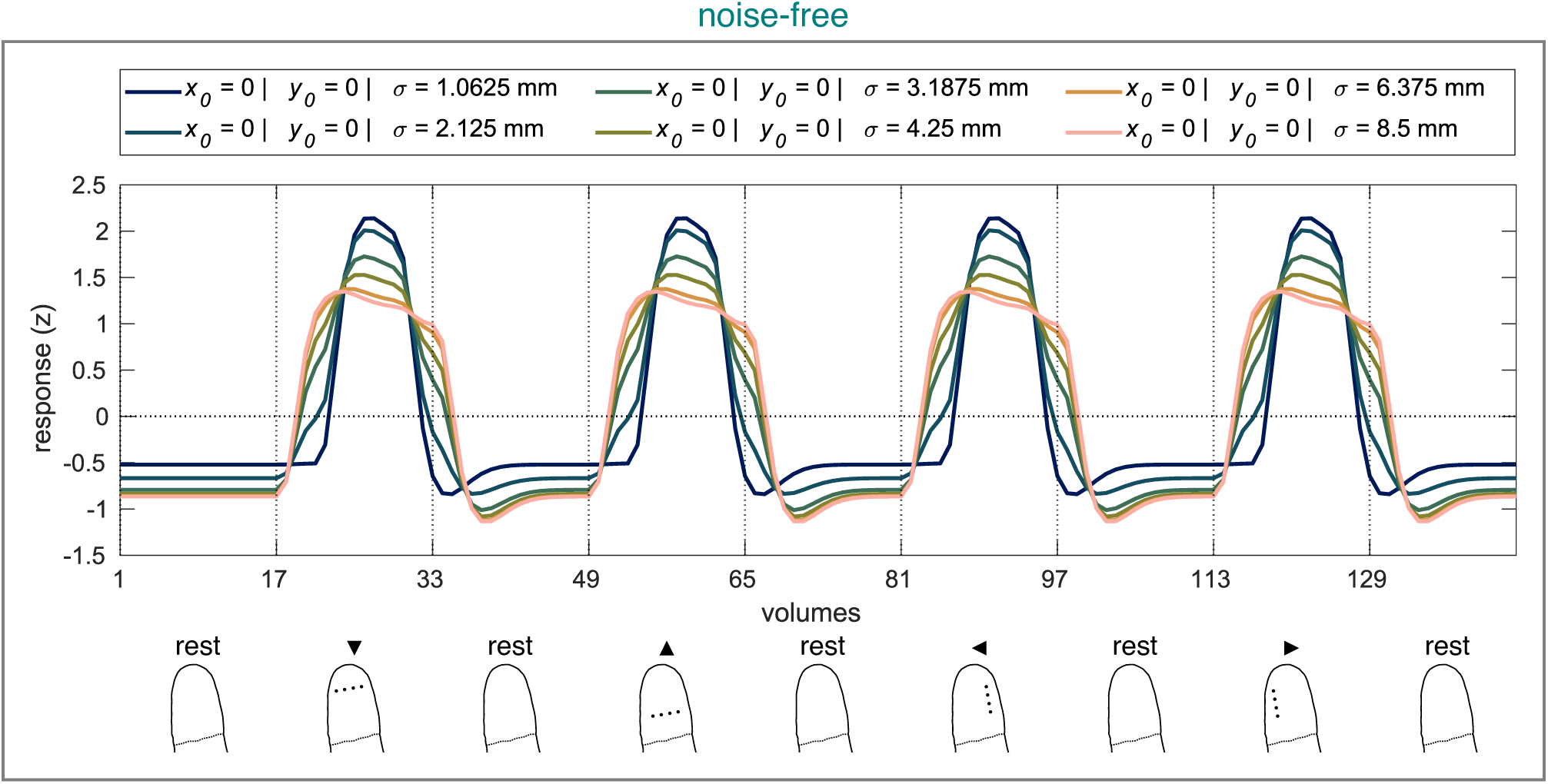
Noise-free time series simulated using the 2dg model with ground-truth pRFs varying in size. Upper panel: Time series corresponding to the ground-truth pRF parameters. Vertical lines accompanied by tick labels denote the onset of stimulation and baseline (rest) intervals. Legend lists ground-truth pRF parameters. Lower panel: Status of the fingertip at the onset of stimulation and baseline (rest) intervals (for more details, see Figure 2). 2dg = 2D Gaussian pRF model. pRF = population receptive field.

**Figure 6.**
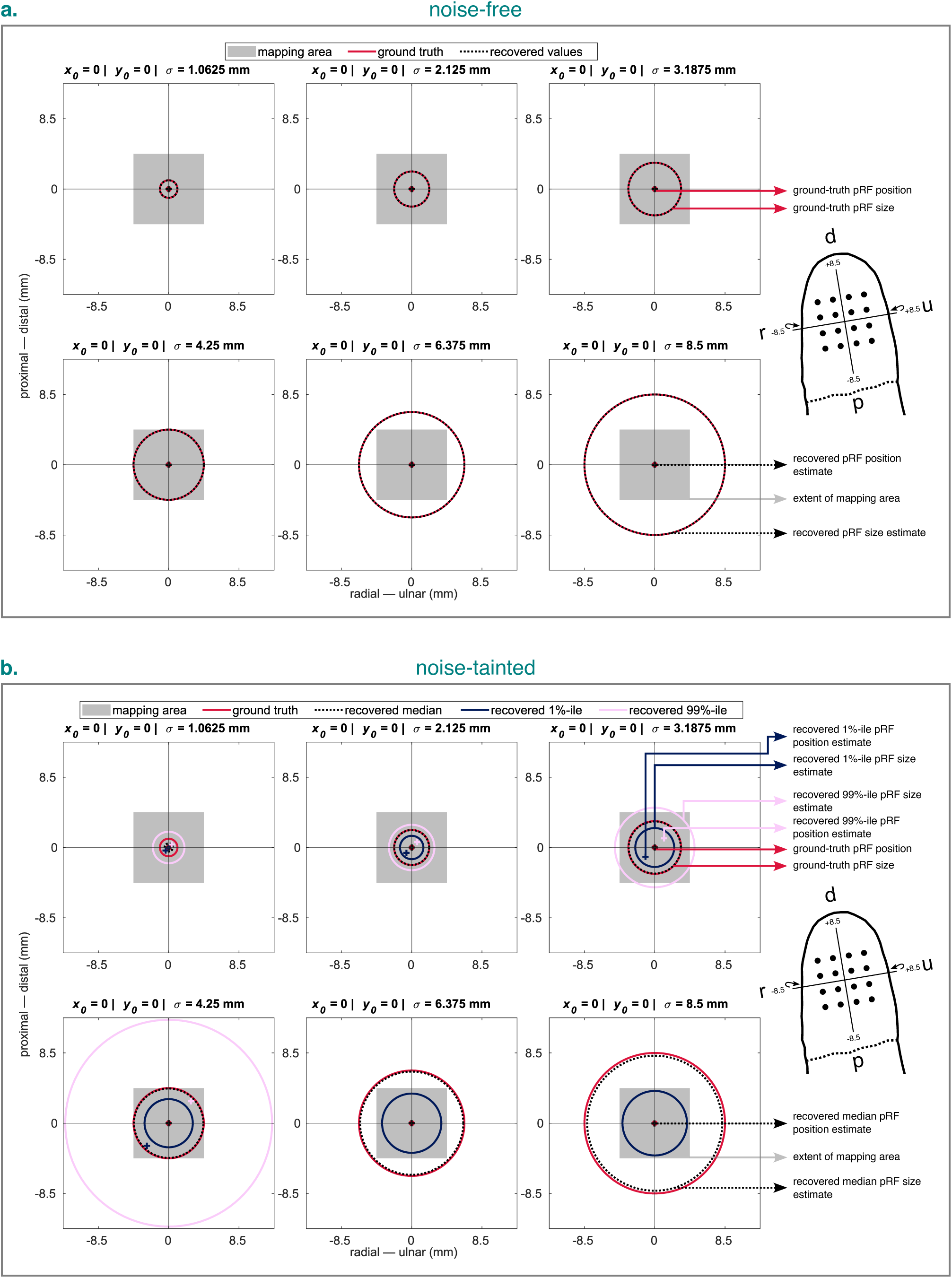
Recovered pRF estimates for noise-free (a.) and noise-tainted (b.) time series simulated and fit using the 2dg model with ground-truth pRFs varying in size. For the noise-tainted time series, 100000 repeats were simulated for each ground-truth pRF. Crosses represent the recovered or ground-truth pRF position. Circles represent the recovered or ground-truth pRF size and are all centered around 0 for better comparison. Note that for the two largest ground-truth pRFs, the lower and upper percentiles for the recovered pRF position estimates and the upper percentile for the recovered pRF size estimate were too large to be displayed. Headers in each panel list ground-truth pRF parameters. Wrap-around arrows indicate an *x*-axis that is slightly wrapped around the fingertip. %-tile = percentile. 2dg = 2D Gaussian pRF model with unconstrained optimization. pRF = population receptive field. d = distal. p = proximal. r = radial. u = ulnar.

Taken together, this pattern of results suggests that for larger pRFs (relative to the mapping area), there is not enough unique information left in a noise-tainted time series to keep the parameter estimates within a plausible range. This does not only lend support to the idea that fingertip pRFs in BA3b are large (relative to the mapping area), but also that we lack the sensitivity to distinguish between different large pRFs, and that an unconstrained optimization procedure with an unlimited parameter space is ill-suited. To address these issues, we therefore constrained the pRF model and fitting procedure. Specifically, we fixed the parameter for pRF size to 8.5 mm, that is, a pRF that basically occupies the whole fingertip, resulting in the 2dg-fix model. We consider such a large pRF size reasonable because research in the macaque and owl monkey indicated that receptive fields of neurons in BA3b can occupy almost the entire fingertip (Merzenich et al., 1978; Nelson et al., 1980; Sur et al., 1980)^5^. In addition to that, we used the constrained grid search fits that are based on a limited parameter space as final pRF estimates. Before applying this refined procedure to our empirical data, though, we performed a series of simulations to investigate its computational validity (accuracy). The corresponding results can be found in S2.1 PRF estimates are computationally accurate within limits and Figure S2-Figure S4.

### 3.3 Fingertip pRFs are organized so that the ulnar-to-radial axis along the fingertip maps onto an inferior-to-posterior axis in BA3b

Figure 7 shows the pRF position estimates (*x*_0_ in a. and *y*_0_ in b.) for the identified BA3b cluster per participant and dataset. The pRF position estimates were obtained using the 2dg-fix model with constrained grid search. Figure S5 displays the corresponding pRF amplitude (*β*_1_ in a.) and pRF baseline (*β*_0_ in b.) parameters. The *x*_0_ position estimates based on all runs indicated that the ulnar-to-radial axis spanning the fingertip maps onto a superior-to-inferior axis in BA3b for both Sub-01 and Sub-02. For Sub-03, however, such a gradient was not clearly visible. A split-half analysis showed that the quantified *x*_0_ map for Sub-01 and Sub-02 was largely reliable across odd and even runs. For Sub-03, however, this analysis flagged that the more inferior part of the *x*_0_ map was rather unreliable. For the *y*_0_ position estimates based on all runs, no clear position gradient was evident across participants with a plethora of estimates being positive and large in magnitude. Nonetheless, just like for the *x*_0_ position estimates, a split-half analysis suggested that the quantified *y*_0_ maps for Sub-01 and Sub-02 were largely reliable across odd and even runs, whereas the inferior part of the *y*_0_ map was rather unreliable for Sub-03.

**Figure 7.**
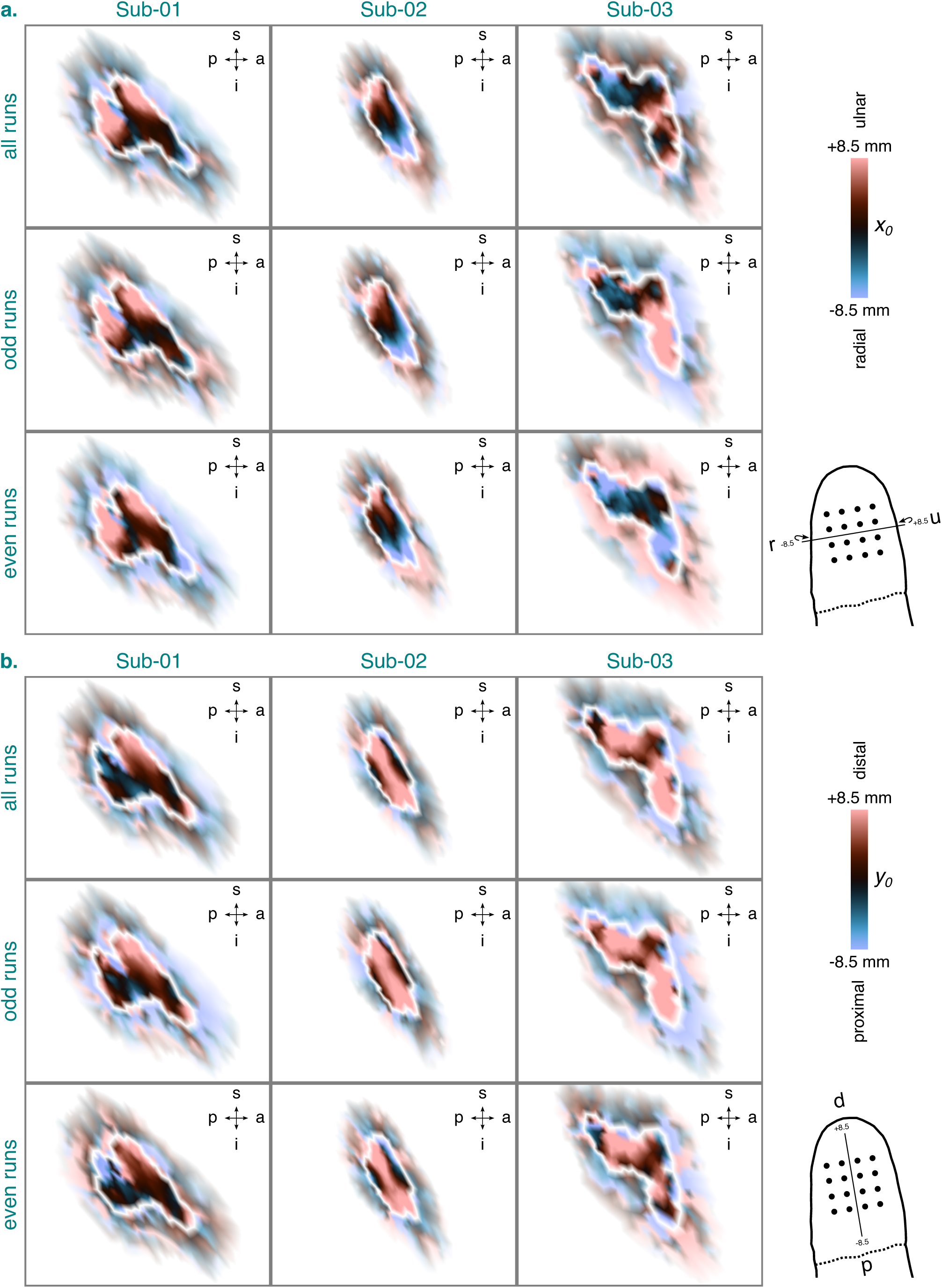
PRF position estimates (*x*_0_ in a. and *y*_0_ in b.) for the identified BA3b cluster and the 2dg-fix model per participant and dataset. The *x*_0_ and *y*_0_ estimates are based on data from session 2-4. They were projected onto an inflated cortical surface model where dark gray regions represent sulci and light gray regions gyri. White lines denote the extent of the identified BA3b cluster (see Figure 4). A zoomed-in view is shown here showing the *x*_0_ and *y*_0_ estimates inside the identified BA3b cluster (opaque) and directly outside of it (semi-transparent). Any *x*_0_ or *y*_0_ estimates surpassing a value of ±8.5 mm were set to this value. Wrap-around arrows indicate an *x*-axis that is slightly wrapped around the fingertip. BA3b = Brodmann area 3b. s = superior. p = posterior. i = inferior. a = anterior. Sub-01, Sub-02, Sub-03 = subject 01, 02, and 03. 2dg-fix = 2D Gaussian pRF model with parameter for pRF size fixed and constrained grid search. pRF = population receptive field. d = distal. p = proximal. r = radial. u = ulnar.

To confirm that the quantified *x*_0_ and *y*_0_ estimates are sensibly connected to the observed data, we inspected the vertex-wise observed and fitted time series for the 2dg-fix model from the identified BA3b cluster ín each participant. Figure 8 illustrates two example vertices for Sub-02 based on all runs. Figure 9 shows the same, but split by odd (a.) and even (b.) runs. When considering all runs, both vertices had a positive *y*_0_ estimate, but opposite-signed *x*_0_ estimates. In line with this, the observed and fitted time series for both vertices showed peaks with a distal bias for intervals stimulating the distal-to-proximal or proximal-to-distal axis along the fingertip, speaking to a positive *y*_0_ estimate. Moreover, for intervals stimulating the ulnar-to-radial or radial-to-ulnar axis, the time series for one vertex showed a peak with a radial bias and the time series for the other vertex a peak with an ulnar bias, speaking to a negative and positive *x*_0_ estimate, respectively. The split-half analysis indicated that these patterns were largely reliable across odd and even runs. Importantly, however, despite the good correspondence between the observed and fitted time series, the fitted time series appeared to be slightly shifted forward in time across all datasets.

**Figure 8.**
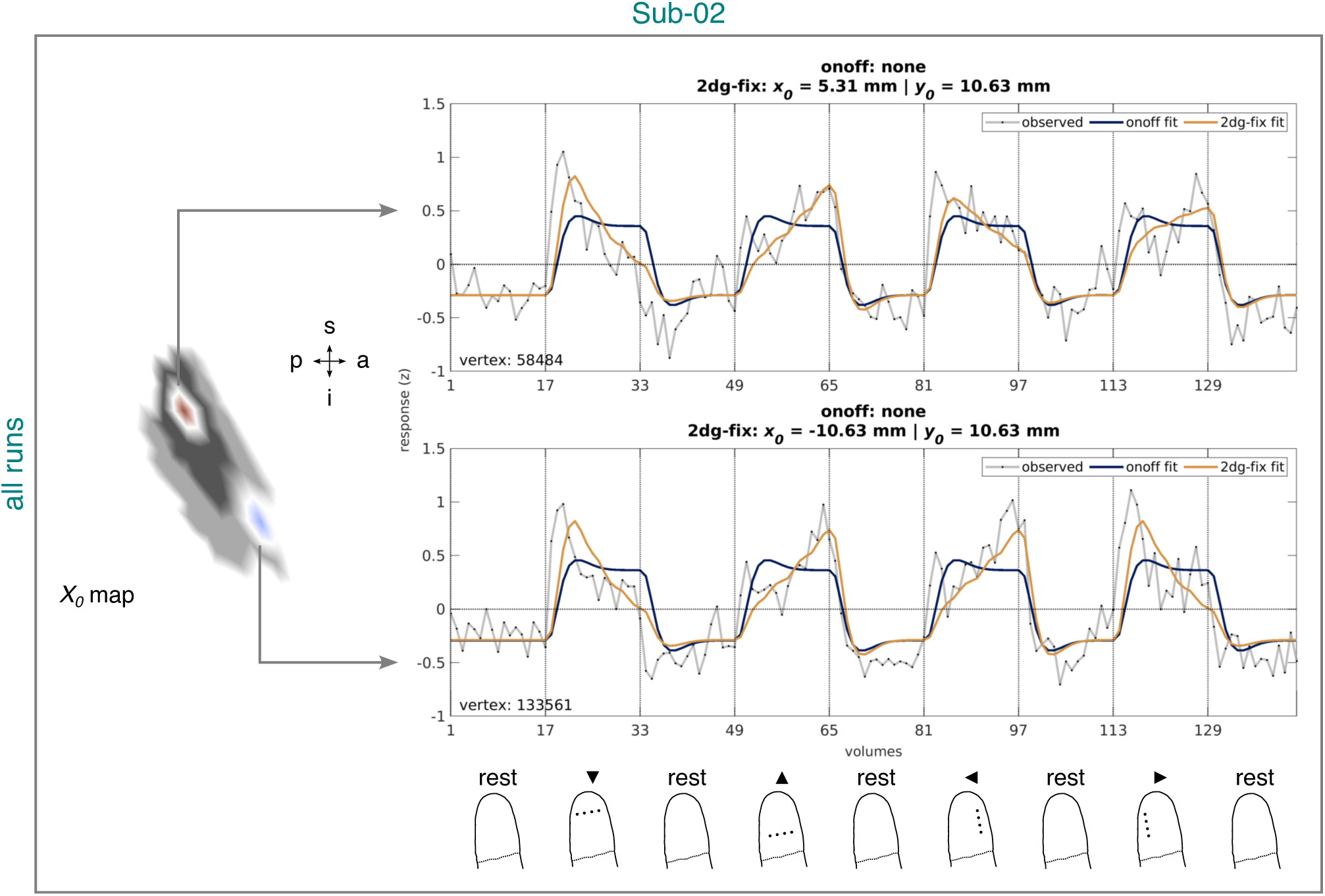
Observed and fitted time series corresponding to the onoff and 2dg-fix model from the identified BA3b cluster for Sub-02 and all runs. The time series are based on data from session 2-4. Left panel: Example vertices from the quantified *x*_0_ map projected onto an inflated cortical surface model where dark gray regions represent sulci and light gray regions gyri (see Figure 7, a.). White lines circumscribe the example vertices. A zoomed-in view is shown here, limited to the extent of the identified BA3b cluster (see Figure 4). Right upper and middle panel: Time series corresponding to the example vertices. Headers in each panel list pRF position estimates (if available) for each model. Vertex identifier is shown on the bottom left of each panel. Vertical lines accompanied by tick labels denote the onset of stimulation and baseline (rest) intervals. Right lower panel: Status of the fingertip at the onset of stimulation and baseline (rest) intervals (for more details, see Figure 2). Sub-02 = Subject 02. 2dg-fix = 2D Gaussian pRF model with parameter for pRF size fixed and constrained grid search. onoff = pRF model without any spatial tuning that simply reflects the presence or absence of tactile stimulation. pRF = population receptive field. BA3b = Brodmann area 3b. s = superior. p = posterior. i = inferior. a = anterior.

**Figure 9.**
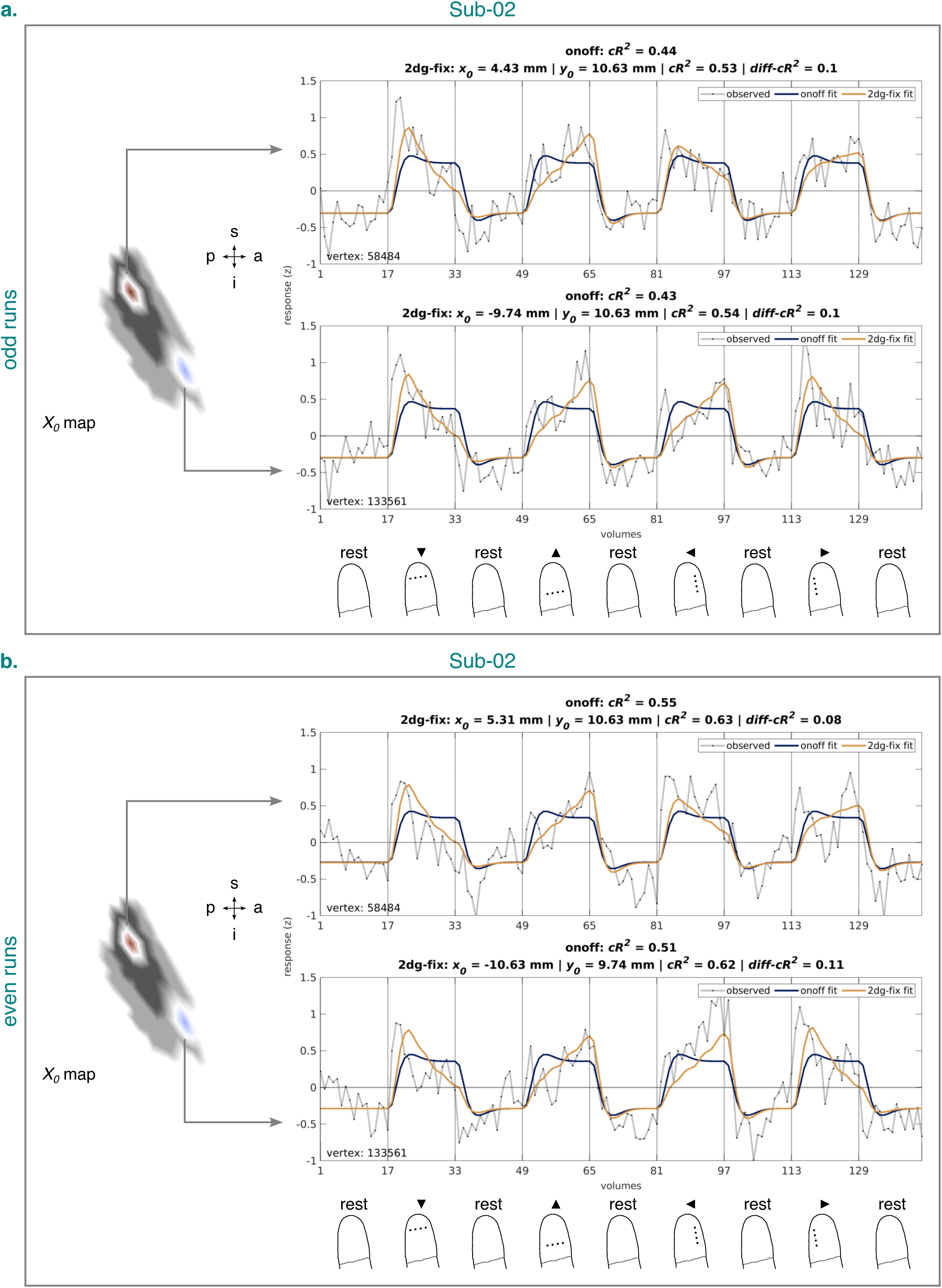
Observed and fitted time series corresponding to the onoff and 2dg-fix model from the identified BA3b cluster for Sub-02 and the odd (a.) and even (b.) runs. The time series are based on data from session 2-4. Left panel (in each subfigure): Example vertices from the quantified *x*_0_ map projected onto an inflated cortical surface model where dark gray regions represent sulci and light gray regions gyri (Figure 7, a.). White lines circumscribe the example vertices. A zoomed-in view is shown here, limited to the extent of the identified BA3b cluster (see Figure 4). Right upper and middle panel (in each subfigure): Time series corresponding to the example vertices. Headers in each panel list pRF position estimates (if available) and cross-validated goodness-of-fit measures for each model. Vertex identifier is shown on the bottom left of each panel. Vertical lines accompanied by tick labels denote the onset of stimulation and baseline (rest) intervals. Right lower panel (in each subfigure): Status of the fingertip at the onset of stimulation and baseline (rest) intervals (for more details, see Figure 2). Sub-02 = Subject 02. 2dg-fix = 2D Gaussian pRF model with parameter for pRF size fixed and constrained grid search. onoff = pRF model without any spatial tuning that simply reflects the presence or absence of tactile stimulation. pRF = population receptive field. BA3b = Brodmann area 3b. s = superior. p = posterior. i = inferior. a = anterior.

Just like for Sub-02, Figure S6-Figure S9 illustrate two example vertices for Sub-01 and Sub-03 based on all runs or split by odd (a.) and even (b.) runs. The time series for the vertices of these participants confirmed the trends we observed for Sub-02. In addition, the time series for the vertices of Sub-01 revealed that more pronounced peaks tended to be associated with *x*_0_ and *y*_0_ estimates of larger magnitude. Moreover, the time series for one vertex of Sub-03 showed that poor signal modulation was linked to unreliable *x*_0_ estimates across odd and even runs. This vertex was located in the inferior portion of Sub-03’s identified BA3b cluster.

In sum, our results speak to a fine-grained somatotopic *x*_0_ position gradient corresponding to the fingertip in BA3b. However, in light of the unclear *y*_0_ position gradient and the imposed modeling constraints, the question arises as to whether we indeed found evidence for spatially-tuned pRFs. To address this question, we next investigated to what extent the 2dg-fix model has superior explanatory power over a model without any spatial tuning (onoff model).

### 3.4 Fingertip pRFs in BA3b are spatially-tuned

Figure 10 shows the cross-validated goodness-of-fit (*cR*^2^) for the 2dg-fix model and the onoff model or the difference thereof (*diff-cR*^2^) for the identified BA3b cluster per participant and cross-validation fold. The cross-validation folds either comprised even runs validated on odd runs (a.) or vice versa (b). Overall, the cross-validated goodness-of-fit for both models was decent for all participants. Moreover, consistent with the split-half analysis for the pRF position estimates, cross-validated goodness-of-fit for the inferior part of Sub-03’s position map was rather low. Importantly, these tendencies appeared to be largely consistent across cross-validation folds. When contrasting the cross-validated goodness-of-fit for the 2dg-fix model and the onoff model directly, the 2dg-fix model slightly outperformed the onoff model in various locations for all participants, which was largely consistent across both cross-validation folds. Specifically, across vertices, participants, and cross-validation folds, the difference in cross-validated goodness-of-fit in favor of the 2dg-fix model ranged from just slightly above 0 to 0.1488^6^ (14.88%). Figure 9, Figure S7, and Figure S9 illustrate the observed and fitted time series corresponding to the onoff and 2dg-fix model from the identified BA3b cluster for each participant. Specifically, two example vertices are shown for the odd (a.) and the even (b.) runs along with the quantified simple (*cR*^2^) and differential (*diff-cR*^2^) values for cross-validated goodness-of-fit. Just like for the pRF position estimates, inspecting these visualizations confirmed that the simple and differential values for cross-validated goodness-of-fit are sensibly connected to the observed data.

**Figure 10.**
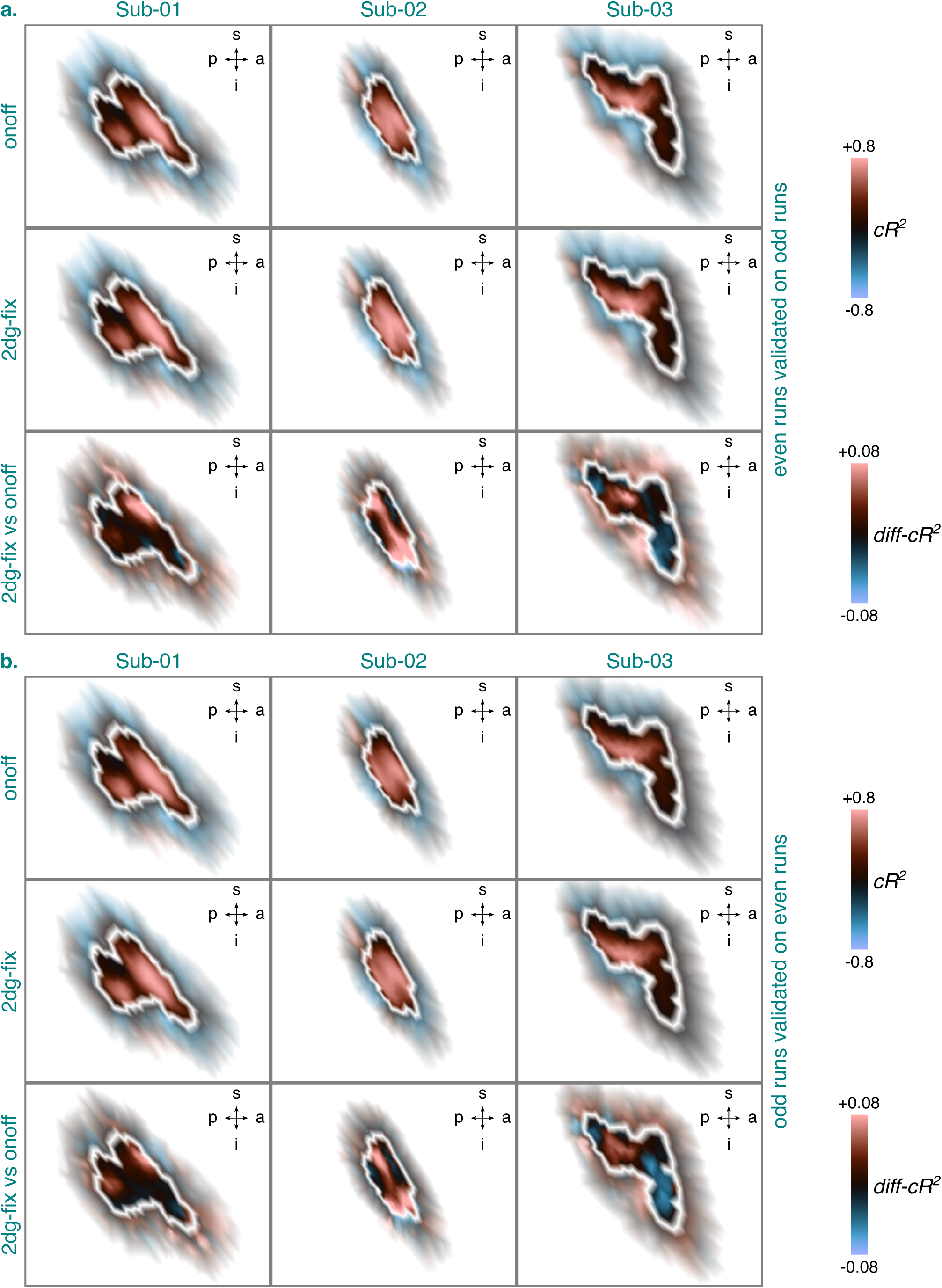
Cross-validated goodness-of-fit (*cR*^2^) for the 2dg-fix and the onoff model or the difference thereof (diff-*cR*^2^) for the identified BA3b cluster per participant and cross-validation fold either consisting of even runs validated on odd runs (a.) or odd runs validated on even runs (b.). The *cR*^2^ and *diff-cR*^2^ values are based on data from session 2-4. They were projected onto an inflated cortical surface model where dark gray regions represent sulci and light gray regions gyri. White lines denote the extent of the identified BA3b cluster (see Figure 4). A zoomed-in view is shown here showing the *cR*^2^ and *diff-cR*^2^ values inside the identified BA3b cluster (opaque) and directly outside of it (semi-transparent). Any *cR*^2^ or *diff-cR*^2^ values surpassing a value of ±0.8 or ±0.08 were set to this value, respectively. BA3b = Brodmann area 3b. s = superior. p = posterior. i = inferior. a = anterior. Sub-01, Sub-02, Sub-03 = subject 01, 02, and 03. 2dg-fix = 2D Gaussian pRF model with parameter for pRF size fixed and constrained grid search. onoff = pRF model without any spatial tuning that simply reflects the presence or absence of tactile stimulation. pRF = population receptive field. *diff-cR*^2^ = differential *cR*^2^.

To investigate whether the quantified differences in cross-validated goodness-of-fit in favor of the 2dg-fix are meaningful and we thus indeed found evidence for spatially-tuned pRFs, we additionally simulated a null distribution. Figure 11 shows the simulated differences in cross-validated goodness-of-fit (*diff-cR*^2^) for the 2dg-fix vs the onoff model per cross-validation fold when noise was added to the simulated data. The cross-validation folds either consisted of even data validated on odd data (a.) or vice versa (b.). The simulated data were obtained using the onoff model and the model fits using the onoff model and the 2dg-fix model with constrained grid search. The difference distribution was clearly negatively skewed. Moreover, the *p*-values calculated for a range of critical differences suggested that even slight differences in favor of the 2dg model are unlikely to be observed by chance and might thus be meaningful. Both these tendencies were consistent across cross-validation folds. Overall, these results not only indicate that we found evidence for spatially-tuned pRFs, but also that the *x*_0_ position gradient we identified appears to be meaningful.

**Figure 11.**
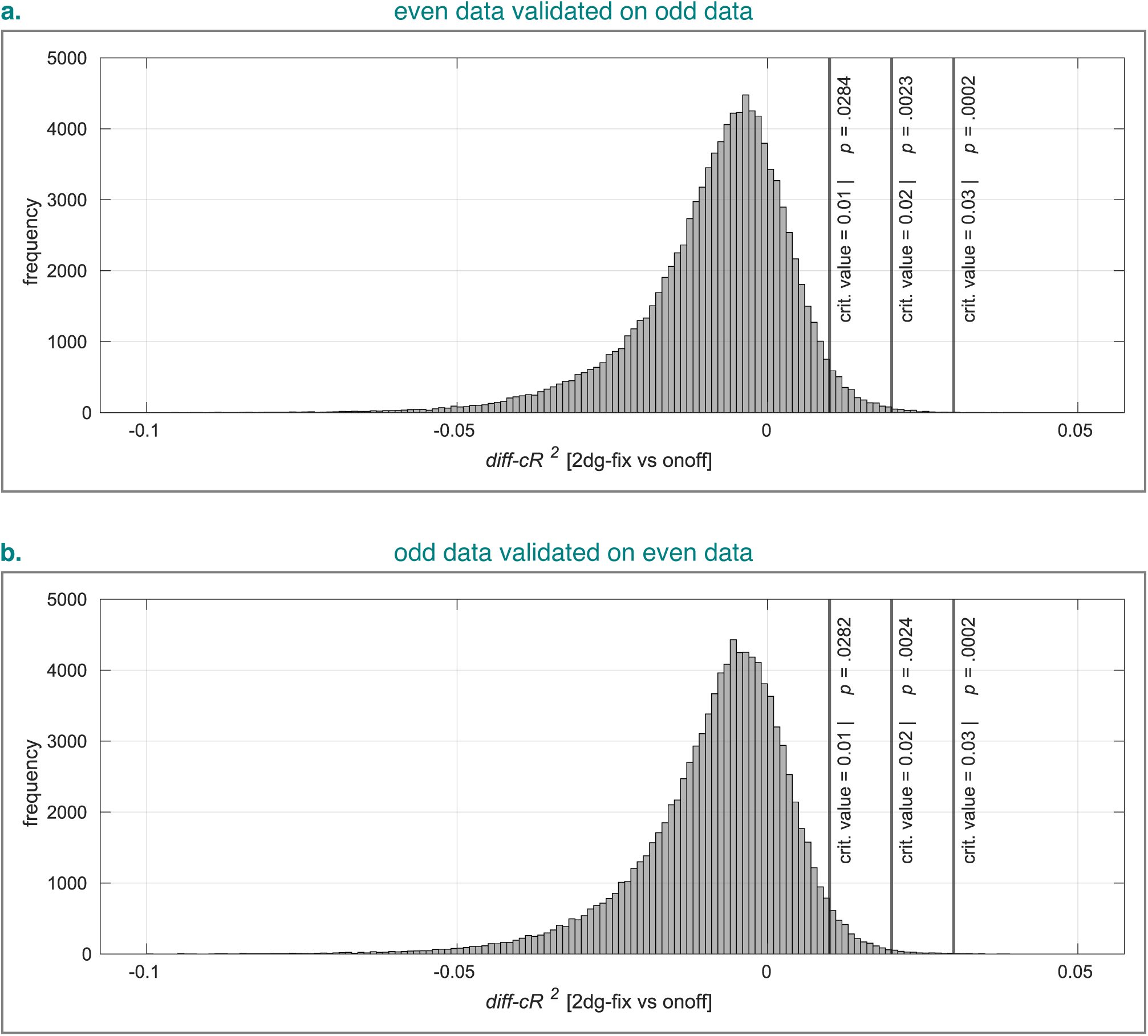
Simulated difference in cross-validated goodness-of-fit for the 2dg-fix vs the onoff model per cross-validation fold either consisting of even data validated on odd data (a) or odd data validated on even data (b). Noise-tainted time series were simulated using the onoff model with 100000 repeats for each cross-validation fold and both the 2dg-fix and the onoff model were fit to the simulated data. Histogram bins ranged from −0.1 to +0.05 with a constant bin width of 0.001. 2dg-fix = 2D Gaussian pRF model with parameter for pRF size fixed and constrained grid search. onoff = pRF model without any spatial tuning that simply reflects the presence or absence of tactile stimulation. pRF = population receptive field. crit. = critical. *diff-cR*^2^ = differential *cR*^2^.

However, any claims about the meaningfulness of observed difference in cross-validated goodness-of-fit require that the simulated null distribution and thus the level of noise we used to perturb noise-free time series was adequate. The same is true for any claims about the computational validity of our pRF estimates. To address these issues, we next investigated to what extent the overall explanatory power, that is, the median cross-validated goodness-of-fit, differs between empirical and simulated data. In doing so, we also investigated to what extent the overall explanatory power varies between participants and fitted models.

### 3.5 Empirical overall explanatory power is variable and similar to simulated one

Table S2 shows the median cross-validated goodness-of-fit (*cR*^2^) for each participant or simulation type as a function of fitted pRF model and cross-validation fold. Whereas the median cross-validated goodness-of-fit varied slightly between cross-validation folds for each participant and fitted pRF model, the variability across participants was much greater, with Sub-02 showing the highest median values, followed by Sub-01 and then by Sub-03. These interindividual differences resonate well with both the split-half analysis for pRF position estimates and prior analyses involving cross-validated goodness-of-fit and suggest that the overall explanatory power was highest for Sub-02. This increase not only needs to be considered when interpreting empirical pRF position gradients, but also provided an upper benchmark for evaluating the appropriateness of our simulations. Considering that the median cross-validated goodness-of-fit for all simulations lay well within the range observed for Sub-02, the overall explanatory power was comparable in this case. As such, the level of noise we used to perturb noise-free time series can be considered adequate.

Apart from these observations, it is worth noting that for all participants except Sub-03, the median cross-validated goodness-of-fit tended to be higher for the fitted 2dg-fix model as compared to the fitted onoff model across both cross-validation folds. Notably, this difference in median cross-validated goodness-of-fit between the fitted models was absent for both cross-validation folds when we simulated data using the onoff model. These findings mirror our previous results for empirical and simulated differences in cross-validated goodness-of-fit, broadly corroborating the idea that we found evidence for spatially-tuned tactile pRFs.

## 4 General discussion

In the present study, we sought to estimate fine-grained 2D spatio-tactile pRFs of the tip of the index finger for the first time in human SI. To this end, we recorded brain responses whilst finely stimulating a portion of participants’ fingertips vibro-tactilely and focused on BA3b as a region of interest. As a first step, we fit a 2D Gaussian pRF model to the data using an unconstrained fitting procedure. This resulted in implausible estimates for pRF position and size, suggesting the recorded brain responses do not contain enough unique information to keep the estimates within a plausible range. Simulations indicated that this likely occurred because the size of pRFs in BA3b exceeds the mapping area on the fingertip skin, resulting in partial pRF mapping and thus less distinct brain responses. As a second step, we therefore constrained the fitting procedure and the 2D Gaussian pRF model by fixing pRF size. When doing so, our results for pRF position indicated that the ulnar-to-radial axis along the fingertip is represented along a superior-to-inferior axis in BA3b, whilst no clear pRF position gradient was discernible for the distal-to-proximal-axis. Although this pattern of results was largely consistent within and across participants (2 out of 3), a cross-validated model comparison showed that our constrained 2D Gaussian pRF model only slightly outperforms a pRF model without any spatial tuning in various cortical locations. However, simulations highlighted that even such slight differences might be meaningful and with that the pRF position estimates we obtained. Collectively, these findings shed first light onto the fine-grained functional architecture of human fingertip maps.

### 4.1 The size of fingertip pRFs in BA3b

Our finding of putatively large fingertip pRFs (relative to the mapping area) in human BA3b appears compatible with invasive recordings in the macaque and owl monkey showing that the area of excitatory receptive fields in BA3b can range from a subportion of to almost the entire fingertip (Merzenich et al., 1978; Nelson et al., 1980; Sur et al., 1980). Although we cannot make any firm statements about the exact size of fingertip pRFs in human BA3b, our simulations suggest that for a pRF sitting at the middle of the mapping area, pRF size is likely larger than the mapping area, and thus 4.25 mm. Such large pRFs can arise from a multitude of neuronal and non-neuronal sources.

#### 4.1.1 Neuronal sources

As far as neuronal sources are concerned, it is important to appreciate that large fingertip pRFs can result from both small receptive fields of single BA3b neurons with a large position scatter or truly large receptive fields with a small position scatter. Similarly, they can arise from only a few large receptive fields of single BA3b neurons biasing the population estimate. Without having an estimate of the variance of receptive field properties within a voxel in BA3b and modeling this aspect explicitly, it is impossible to distinguish between these possibilities.

Leaving these ambiguities aside, it is conceivable that large fingertip pRFs reflect the existence of receptive fields elongated along the distal-to-proximal axis of the finger. In line with this, invasive recordings in macaques and owl monkeys show that the shape of receptive fields in BA3b corresponding to the fingertips can range from circular to elliptical (Merzenich et al., 1978; Nelson et al., 1980; Sur et al., 1980). Human fMRI studies modeling pRFs along the within- and between-finger dimension furthermore suggest that pRF size in BA3b is larger for the within-finger dimension (although the exact physical interpretation of this finding remains unclear, as the modeling was performed in abstract sensory space; Asghar et al., 2023; Wang et al., 2021). As such, expanding the mapping area along the distal-to-proximal axis of the finger might be critical for obtaining a direct estimate of the size and eventually also the shape of fingertip pRFs.

Similarly, it is possible that large fingertip pRFs evince the existence of multi-digit receptive fields. In fact, electrophysiological studies in macaques suggest that a subportion of BA3b neurons responds to tactile stimulation to multiple fingers (DiCarlo et al., 1998; Lazar et al., 2022; Trzcinski et al., 2023). Human fMRI studies modeling pRFs along the between-finger(tip) dimension, sometimes in combination with the within-finger dimension, also show tendencies for multi-digit pRFs, although it is not always clear whether these are indeed present in BA3b (Asghar et al., 2023; P. Liu et al., 2021; Puckett et al., 2020; Schellekens et al., 2021; Wang et al., 2021). Importantly, these considerations suggest that it might be necessary to expand the mapping area to other fingers to obtain comprehensive estimates of the size as well as shape of fingertip pRFs.

Lastly, large fingertip pRFs might be related to the type of stimulation we applied. Specifically, we adopted a position-changing vibrotactile stimulus consisting of a row of pins that oscillated at a frequency of 25 Hz with brief on and off periods. This stimulus might have not only been effective in driving Merkel cell neurite complexes and Meissner corpuscles, mechanoreceptors known to have small peripheral receptive fields (area: ∼9 mm^2^ and ∼22 mm^2^, respectively), but also Pacinian corpuscles, mechanoreceptors known to have large peripheral receptive fields (e.g., entire finger; as summarized in Johnson, 2002). This is because all these mechanoreceptors are known to be sensitive to a range of frequencies including 25 Hz (Merkel: ∼0-100 Hz; Meissner: ∼1-300 Hz; Pacinian: ∼5-1000 Hz), albeit their peak sensitivities are thought to differ markedly (Merkel: ∼5 Hz; Meissner: ∼50 Hz; Pacinian: ∼200 Hz; as summarized in Johnson, 2002). What is more, locally applied vibrotactile stimulation is known to result in surface waves propagating through the skin, effectively enlarging the stimulus and limiting its resolution (Dandu et al., 2021; Manfredi et al., 2012; Sofia & Jones, 2013; Tummala et al., 2024). Depending on how pronounced and far-reaching these propagating waves were, they might have led to a fairly effective stimulation of the large peripheral receptive fields of Pacinian corpuscles despite our small mapping area, which might have manifested itself at the cortical level too.

#### 4.1.2 Non-neuronal sources

As far as non-neuronal sources are concerned, it may be argued that large pRF sizes are due to the brain’s macrovasculature. This is because fMRI measures a hemodynamic response (Logothetis, 2003) and GE BOLD fMRI, as applied in our study, is known to be sensitive to large draining veins, such as those near the pial surface (Bause et al., 2020; Koopmans et al., 2010; Polimeni et al., 2010). This sensitivity might lead to a systematic blurring of fMRI time series near the pial surface and thus an inflation of pRF size estimates. It is important to note, however, that our analyses were limited to voxels located midway between vertices on the gray-white matter boundary and vertices on the pial boundary. Moreover, pRF size estimates in human visual cortex based on GE BOLD-fMRI have been shown to vary according to a u-shaped function across cortical depth (Fracasso et al., 2016). And lastly, visual pRF mapping in macaques shows that GE BOLD-fMRI-based pRF size estimates and variations thereof closely reflect those obtained via direct neural recordings (Klink et al., 2021).

As such, our finding of putatively large pRF sizes (relative to the mapping area) cannot be easily dismissed as a mere effect of the brain’s macrovasculature.

Another factor that may have led to a blurring of the fMRI time series and thus an inflation of pRF size estimates is body motion, such as movements of the head or hand. However, the effect of body motion on fMRI time series is likely complex, depending, amongst other things, on the nature and location of movement, how body movements interact with the image acquisition process (e.g., T. T. Liu, 2017; Zaitsev et al., 2015), and how and if cutaneous, proprioceptive, and motor information is integrated in BA3b (e.g., Kim et al., 2015). As such, a one-to-one relationship between an increase in body motion and an increase in pRF size seems unlikely. Moreover, it is important to consider that 2 of our 3 participants were highly experienced fMRI participants and all participants familiarized themselves with the pRF mapping experiment prior to scanning, had their hand stabilized using a cushion, and were explicitly instructed to keep still. In addition, our data were prospectively corrected for within-run head motion and retrospectively for between-run head motion via a carefully designed procedure. As such, it seems unlikely that hand or head motion was a primary driving factor.

### 4.2 The organization of fingertip pRFs in BA3b

Our finding of a pRF position gradient where the ulnar-to-radial axis along the fingertip maps onto a superior-to-inferior axis in BA3b fits in with invasive recordings in the macaque and owl monkey showing a similar topographic organization (Merzenich et al., 1978; Nelson et al., 1980). However, unlike these invasive recordings, our findings revealed no clear pRF position gradient for the distal-to-proximal axis along the finger(tip) in BA3b. Similar to our incapacity to obtain a direct estimate of pRF size, this might be due to large receptive fields that are elongated along the distal-to-proximal axis (Merzenich et al., 1978; Nelson et al., 1980; Sur et al., 1980) and a mapping area that was too small to account for this.

Importantly, the pRF position gradient we uncovered for the ulnar-to-radial axis was not discernible for 1 out of our 3 participants. Given that the inferior part of both pRF position maps was rather unreliable for this participant and characterized by a low cross-validated goodness-of-fit, this inter-individual variability seems, at least partly, due to an increased level of noise for this participant. Importantly, however, invasive recordings in owl monkeys and squirrel monkeys point to considerable inter-individual variability in the architecture of hand maps in BA3b (Merzenich et al., 1987). It is therefore vital to assess the test-retest reliability as well as cross-validated goodness-of-fit within a given participant and across participants to distinguish intra-from inter-individual variation in pRF properties, as we did here.

### 4.3 Study limitations

PRF modeling is only as sensible as the data and assumptions it operates upon. To collect high-quality datasets, we combined 7T-fMRI at submillimeter resolution, precision neuroscience, PMC, and fine-grained vibrotactile stimulation. Despite this effort, our findings suggest that the usefulness of these datasets is limited because of partial pRF mapping. Partial pRF mapping is a recognized phenomenon in vision science, especially in higher-level visual areas where pRF size tends to be large (e.g., Mackey & Curtis, 2017; Mackey et al., 2017). Importantly, however, unlike our tactile pRF mapping stimulus, visual pRF mapping stimuli typically sample sensory space more densely and without gaps. As such, the informational value of our fMRI time series is likely much lower. Future studies aimed at resolving fine-grained 2D spatio-tactile pRFs should therefore strive to map out body parts as completely as possible whilst ensuring a dense sampling of the skin. Yet, due to the general lack of commercially-available large-scale and high-density tactile stimulation equipment that is MRI-compatible, this is a challenging endeavor.

Besides, both the explanatory power and computational accuracy of our pRF estimates may be limited by the assumptions we made about the HRF in BA3b. Given that we adopted a canonical HRF, we assumed invariance across voxels(vertices) and individuals, which may not have been the case. Moreover, we assumed that this canonical HRF optimally approximates the properties of empirically-observed data in BA3b. However, this may have only been partially true. Specifically, the slight temporal mismatch between the observed and the fitted fMRI time series we detected for all of our participants may indicate that the canonical HRF was slightly suboptimal. However, accurately estimating ROI-specific, voxel-wise(vertex-wise) HRF parameters in individual brains is challenging and susceptible to noise, thus likely requiring a large amount of additional data.

The explanatory power and computational accuracy may be furthermore compromised by the assumptions we made about the resolution and extent of tactile stimulation. More precisely, although vibrotactile stimulation is known to result in surface waves propagating through the skin (Dandu et al., 2021; Manfredi et al., 2012; Sofia & Jones, 2013; Tummala et al., 2024), the way we represented the tactile stimulus as part of our pRF modeling pipeline does not account for this. Before any such incorporation can happen, though, it is necessary to quantify the properties of propagating waves for a given pRF mapping design via suitable techniques, such as vibrometry (Dandu et al., 2021; Manfredi et al., 2012; Tummala et al., 2024).

Lastly, the adequateness of our simulations might be limited by the assumptions we made about the noise. Specifically, we assumed additive white Gaussian noise across voxels(vertices). As such, we did, for instance, not account for potential noise correlations between neighboring voxels(vertices), potential differences in the level of noise across voxels(vertices), or different sources of noise (e.g., T. T. Liu, 2017; Welvaert & Rosseel, 2014; Zhang et al., 2020). Pitting different noise models against one another is beyond the scope of the present article, but represents an important avenue for future research.

### 4.4 Alternative pRF models and fitting strategies

To address the limited informational value of our fMRI time series when fitting a 2D excitatory Gaussian pRF model, we fixed the parameter for pRF size and let go of unconstrained optimization, relying solely on constrained grid search. This is of course only one way of approaching this problem. There are numerous other approaches, some of which we briefly describe here.

One alternative approach could involve exploring other 2D pRF models that have been previously adopted in vision science, such as a 2D excitatory anisotropic Gaussian pRF model (e.g., Lerma-Usabiaga et al., 2021), a 2D excitatory-inhibitory difference of Gaussians pRF model (e.g., Zuiderbaan et al., 2012), or a 2D excitatory Gaussian pRF model with compressive spatial summation (e.g., Kay et al., 2013). However, given that these alternative pRF models can be regarded as more complex versions of a 2D excitatory Gaussian pRF model, it is unclear why unconstrained optimization would produce plausible pRF estimates for them, but not for a simpler model. Moreover, these alternative pRF models introduce additional free parameters, complicating the fitting procedure and demanding informative fMRI time series. Relatedly, given that our stimulus was sparse and presumably covered only a small portion of the pRFs in BA3b, model differentiation likely becomes difficult.

A second alternative approach might be to use a constrained optimizer with bounds on the parameters for pRF position and pRF size or a set of equations these parameters need to satisfy. A third alternative approach might be to still use an unconstrained optimizer, but add penalties, and thus a cost if the parameter values are too extreme (instead of having hard constraints). As compared to our approach, constrained optimization might work less well as the optimization can still stop anywhere inside the bounds, which might produce unstable, arbitrary solutions. Similarly, penalized optimization might work less well as the penalty might dictate the solution without reflecting the data much. Put differently, given that optimizers receive very little guidance from the data in our study, it might be challenging to use them.

A fourth alternative approach could consist of using the *β* estimates from a GLM as the basis for pRF modeling instead of fMRI time series (e.g., Centanino et al., 2024; Kay et al., 2013; Prince et al., 2022). However, this way, the fitting would be based on fewer data points and the temporal information discarded. In situations where the informational value of fMRI time series is low, this might make it harder to obtain accurate pRF estimates and tease out subtle differences between a spatially-untuned and a spatially-tuned pRF model. In any event, future research is necessary to shed light onto the potential of alternative pRF models and fitting strategies to capture the fine-grained organization of human fingertip maps in BA3b based on our datasets.

## 5 Summary and conclusion

Despite the great importance of fingertip sensations for everyday life, little is known about how tactile input to the fingertips is represented in human SI. Using 2D spatio-tactile pRF modeling, we show – for the first time – that fingertip pRFs in human BA3b may be large (relative to the mapping area) and organized so that the ulnar-to-radial axis across the fingertip maps onto a superior-to-inferior axis in human BA3b. Although we were unable to obtain a direct estimate of pRF size and a clear pRF position gradient for the distal-to-proximal axis along the fingertip, our results bring us closer to understanding the fine-grained architecture of fingertip maps in the human brain.

## Data and code availability

Custom code associated with this manuscript is available via our GitHub repository (Stoll et al., 2026a). Data associated with this manuscript including data produced by executing the custom code are available via our OpenNeuro repository (Stoll et al., 2026c). These repositories are linked components of our OSF repository (Stoll et al., 2026b) that contains general instructions for repository usage and preparatory steps (see README.md). The OSF repository also hosts additional materials, such as the figures and videos presented in this manuscript.

## Contributions

SS: Conceptualization, Methodology, Software, Validation, Formal analysis, Investigation, Data curation, Writing – Original Draft, Writing – Review & Editing, Visualization, Project administration. FL: Methodology, Software, Validation, Formal analysis, Investigation, Data curation, Writing – Review & Editing. DSS: Conceptualization, Methodology, Software, Validation, Data curation, Writing – Review & Editing. HM: Methodology, Investigation, Writing – Review & Editing. PL: Investigation, Writing – Review & Editing. JN: Investigation, Writing – Review & Editing. EK: Conceptualization, Resources, Writing – Review & Editing, Supervision, Project administration, Funding acquisition.

## Acknowledgments

The present work received funding from the European Structure and Investment Fund (ZS/2020/05/ 141591), the European Research Council under the European Union’s Horizon 2020 research and innovation programme (949609) as well as the Deutsche Forschungsgemeinschaft (MA, 9235/3-1, 501214112 and KU 3711/4-1, 528318902). We thank Astrid Wollrab and Cindy Lübeck for help with data collection, Juliane Döhler for discussions about the quality of our segmentation results, Udo Adler for computing support, and the University Clinic for Oral and Maxillofacial Surgery in Magdeburg for manufacturing high-quality mouthpieces..

## Supplementary material S1

### S1 Methods

#### S1.1 Preprocessing

##### Explanation S1

For skull-stripping, we adopted a deep-learning strategy because this strategy is known to be accurate and robust (Hoopes et al., 2022).

##### Explanation S2

For generating templates as well as for alignment and coregistratiom purposes, we adopted nearest neighbor interpolation to keep spatial blurring minimal.

##### Explanation S3

Due to non-brain tissue, alignment issues can arise when generating an unmasked anatomical template across sessions directly. To counteract such issues, we adopted a two-step procedure where we first generated a masked anatomical template and then used the resulting transformation matrices to generate an unmasked anatomical template.

##### Explanation S4

The final masked anatomical template was generated because masking a template (instead of generating it from already masked images) should be more precise.

##### Explanation S5

For anatomical cortical surface reconstruction, we used an anatomical template. This is because the usage of an anatomical template (with a reduced level of noise due to data aggregation; Lüsebrink et al., 2017, 2021) instead of a single anatomical image is known to facilitate the construction of the surface model for anatomical data at submillimeter resolution (Tian, Bilgic, et al., 2021; Tian, Zaretskaya, et al., 2021).

##### Explanation S6

We coregistered the final masked anatomical template (whole-brain data) to the masked functional template (partial-brain data) because it is easier to register whole-brain data to partial-brain data (instead of vice versa), especially when non-brain tissue is being removed.

##### Explanation S7

We applied multiple transformation matrices in one go because it reduces the number of interpolation steps and thus potential interpolation artifacts.

##### Explanation S8

BBR aligns a cortical surface reconstruction (based on a high-quality image with good anatomical contrast) to the gray-white matter boundary in the functional data (with a rather minimal amount of gray-white contrast; Greve & Fischl, 2009; Polimeni et al., 2018). As such, it is well-suited for surface-based analyses relying on accurate positioning of the cortical surface model relative to the functional data (Polimeni et al., 2018). It is also known to excel at aligning partial-brain functional data to whole-brain anatomical data (Greve & Fischl, 2009). This is exactly our use case.

##### Explanation S9

We used the functional voxel located midway between vertices on the gray-white matter boundary and the pial boundary to reduce partial volume effects and the effect of large draining veins residing on the pial surface (Bause et al., 2020; Koopmans et al., 2010; Polimeni et al., 2010).

##### Explanation S10

We modestly smoothed the functional data along the cortical surface to reduce noise.

### S1.2 Data analysis

#### S1.2.1 Simulations

##### Computational validity

To computationally validate the 2dg-fix model with constrained grid search, we simulated how well we can recover the parameters of ground-truth pRFs varying in center position when we add or do not add noise to the data. Whereas the latter allows us to identify flaws in our adapted pRF fitting pipeline, the former permits us to approximate the accuracy of empirical pRF estimates. To this end, we first used the 2dg-fix model to simulate noise-free time series for pRFs with 25 different *x*_0_ positions (from −10.625 to +10.625 mm with 25 evenly-spaced values) and 25 different *y*_0_ positions (from −10.625 to +10.625 mm with 25 evenly-spaced values), matching all 625 search grid value pairs (see 2.7.3 PRF modeling and Figure S2, a., left panel). We then selected a subset of these time series (i.e., the 4*^th^*, 8*^th^*, 12*^th^*, 16*^th^* and 20*^th^* value out of the 25 *x*_0_ and *y*_0_ positions, respectively), reflecting 25 different search grid value pairs (see Figure S3, upper panel), and perturbed each of these time series 100000 times with random Gaussian noise (*σ_noise_* = 2, expressed in units of mean percent overlap with the pRF model). We did this twice to simulate an odd and even dataset. Next, we fit the 2dg-fix model with constrained grid search to both the simulated noise-free and noise-tainted time series of each dataset. In the noise-tainted case, we then determined the number of times (out of 100000) the constrained grid search resulted in a specific grid search value pair for a given ground-truth pRF, separately for each dataset.

Subsequently, we repeated this analysis for ground-truth pRF positions falling halfway between neighboring search grid value pairs. In the noise-free case, we simulated time series for pRFs with 24 different *x*_0_ positions (from −10.625 to +10.625 mm with 49 evenly-spaced values whilst retaining every second value) and 24 different *y*_0_ positions (from −10.625 to +10.625 mm with 49 evenly-spaced values whilst retaining every second value), reflecting 576 locations not matching any search grid value pairs (see Figure S2, b., left panel). In the noise-tainted case, we again selected a subset of these time series (i.e., the 4*^th^*, 8*^th^*, 12*^th^*, 16*^th^* and 20*^th^* value out of the 24 *x*_0_ and *y*_0_ positions, respectively), resulting in 25 locations not matching any grid search value pairs (see Figure S4, upper panel). Note that selecting ground-truth pRF positions that coincide with the search grid value pairs should result in 100% accuracy for the noise-free case assuming our pipeline is unbiased. Due to the discretizing nature of grid search fits, an accuracy of 100% is, however, not achievable for ground-truth pRF positions not matching the search grid value pairs in the noise-free case. Lastly, just like before, we determined the median cross-validated goodness-of-fit for all noise-tainted simulations (see 2.7.3 PRF modeling and 2.7.4 Simulations).

## S2 Results

### S2.1 PRF estimates are computationally accurate within limits

Figure S2 (a.) shows the recovered pRF parameters (*x*_0_, *y*_0_) for ground-truth pRFs varying in position and coinciding with the search grid value pairs when no noise was added to the simulated data. The simulated data were obtained using the 2dg-fix model and the model fits via the 2dg-fix model with constrained grid search. For all simulated pRF positions, we were able to recover the ground truth with an accuracy of 100%. Figure S3 shows the frequency of recovered pRF parameters for the same scenario when noise was added to the simulated data, albeit only for a subset of ground-truth pRF positions. For almost all simulated pRF positions, the most frequently recovered pRF parameters were solely located in close vicinity of the ground truth. For simulated pRF positions close to the boundary of the search space, however, the most frequently recovered pRF parameters were additionally located at the very edge of the search space. Such edge artifacts are expected due to the limits of the search space. Importantly, for all simulated pRF positions, the less frequently recovered pRF parameters were considerably scattered around the ground truth.

To assess the generality of these results, we repeated the simulation analysis for ground-truth pRFs whose position did not coincide with the search grid value pairs. Figure S2 (b.) shows the corresponding recovered pRF parameters (*x*_0_, *y*_0_) for the noise-free case and Figure S4 for the noise-tainted case. Whereas the noise-tainted case largely mirrored our previous observations, the noise-free case additionally illustrated that for all simulated pRF positions, the recovered pRF parameters are systematically biased, albeit still very close to the ground truth. Such biases are expected due to the discretized nature of the search space.

Taken together, these results illustrate that under conditions of no noise, our pRF estimates are computationally accurate, but might suffer from small systematic biases induced by the discretization of the search space. Under conditions of noise, the computational accuracy is further compromised by edge artifacts and a non-negligible spatial uncertainty of parameter estimates. It is important to keep these aspects in mind when interpreting any empirical pRF position gradients.

## Supplementary figures

**Figure S1.**
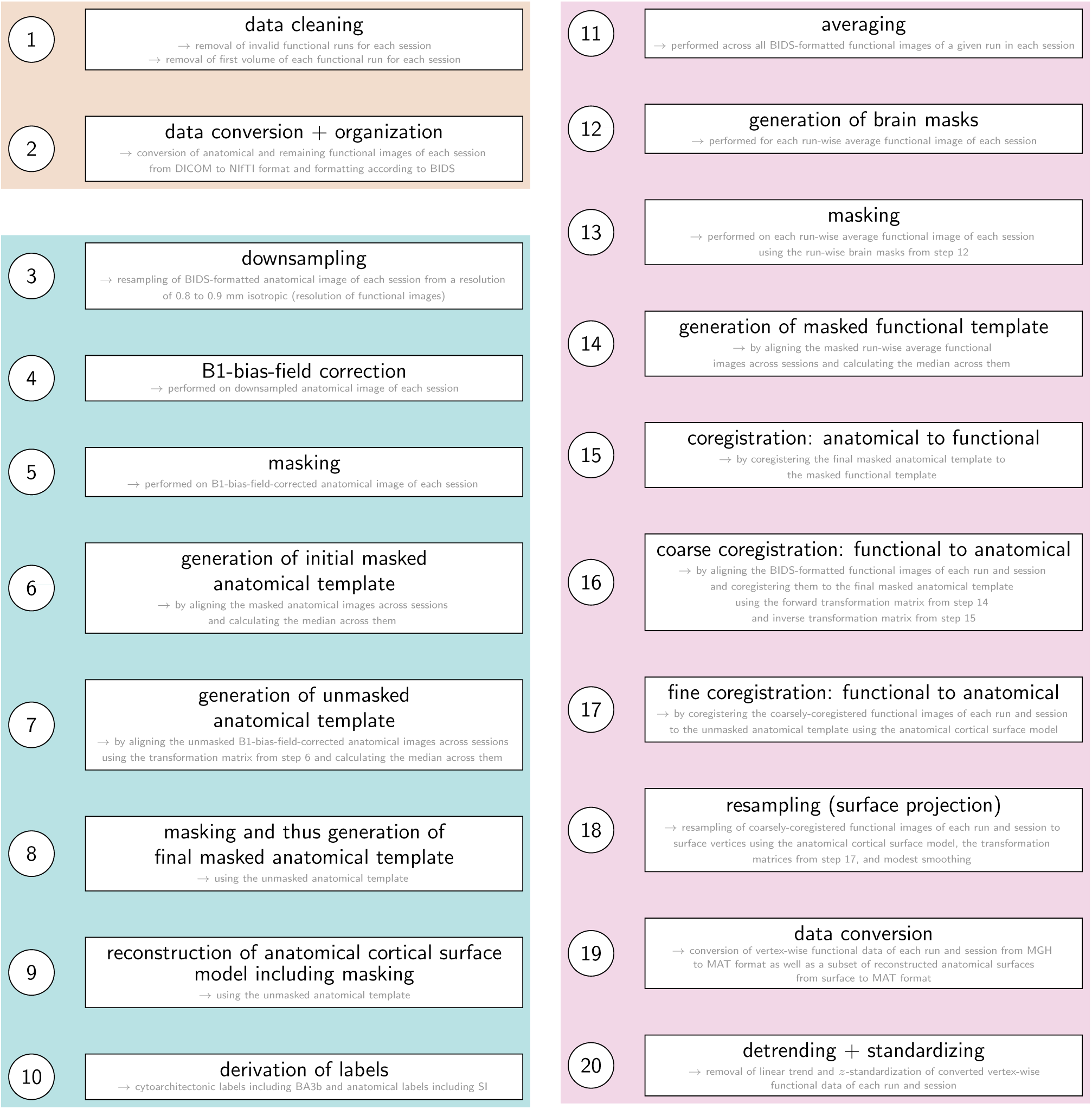
Overview of performed preprocessing steps. Tan box: Preparatory steps. Teal blue box: Preprocessing of anatomical data. Thistle box: Preprocessing of functional data. BA3b = Brodmann area 3b. SI = primary somatosensory cortex. BIDS = Brain Imaging Data Structure.

**Figure S2.**
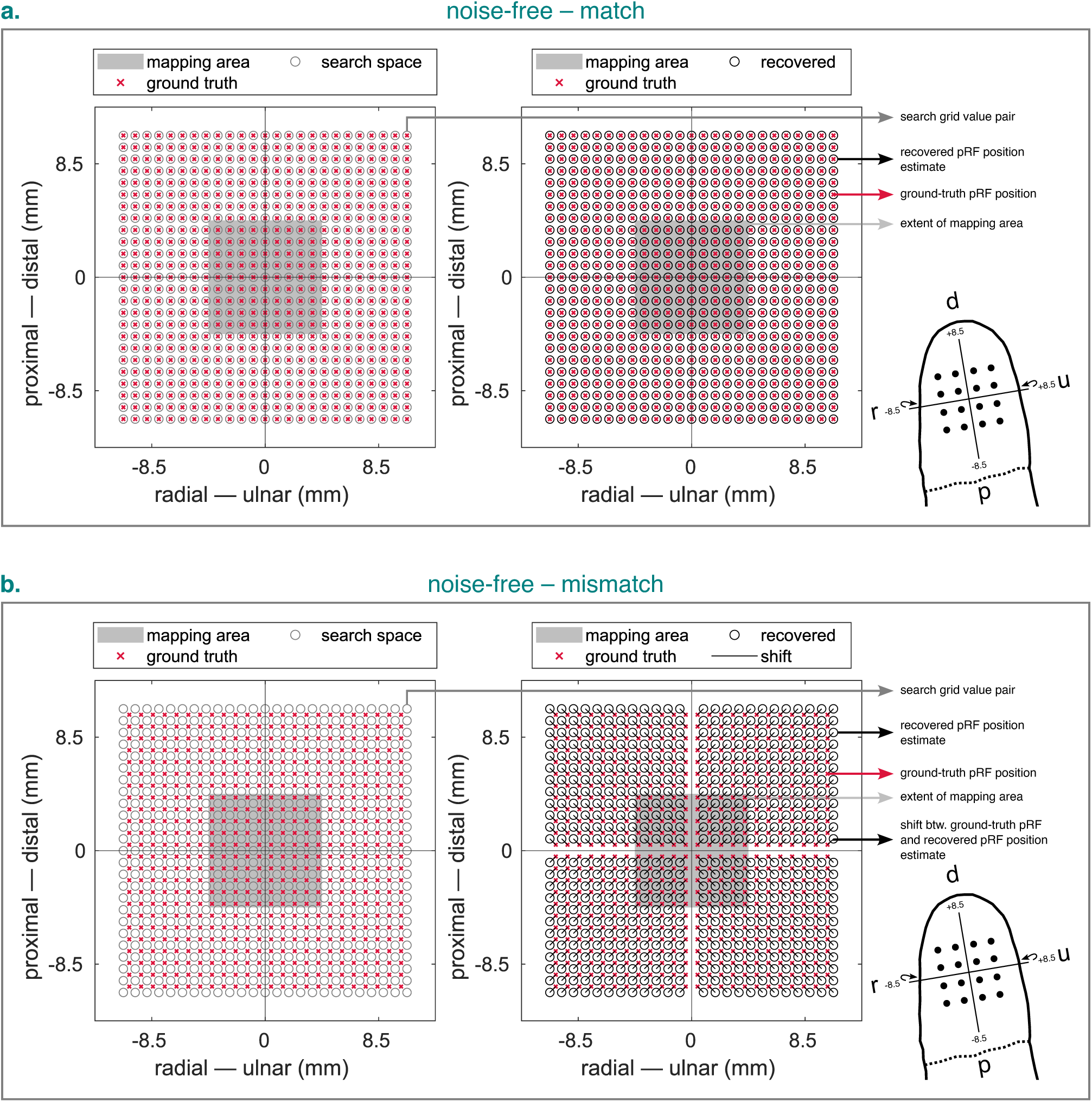
Recovered pRF position estimates for noise-free time series simulated and fit using the 2dg-fix model with ground-truth pRFs matching (a.) or mismatching (b.) the search grid value pairs. Left panel (in each subfigure): Ground-truth pRF positions relative to search grid value pairs. Right panel (in each subfigure): Ground-truth pRF positions relative to recovered pRF position estimates. Wrap-around arrows indicate an *x*-axis that is slightly wrapped around the fingertip. 2dg-fix = 2D Gaussian pRF model with parameter for pRF size fixed and constrained grid search. pRF = population receptive field. d = distal. p = proximal. r = radial. u = ulnar.

**Figure S3.**
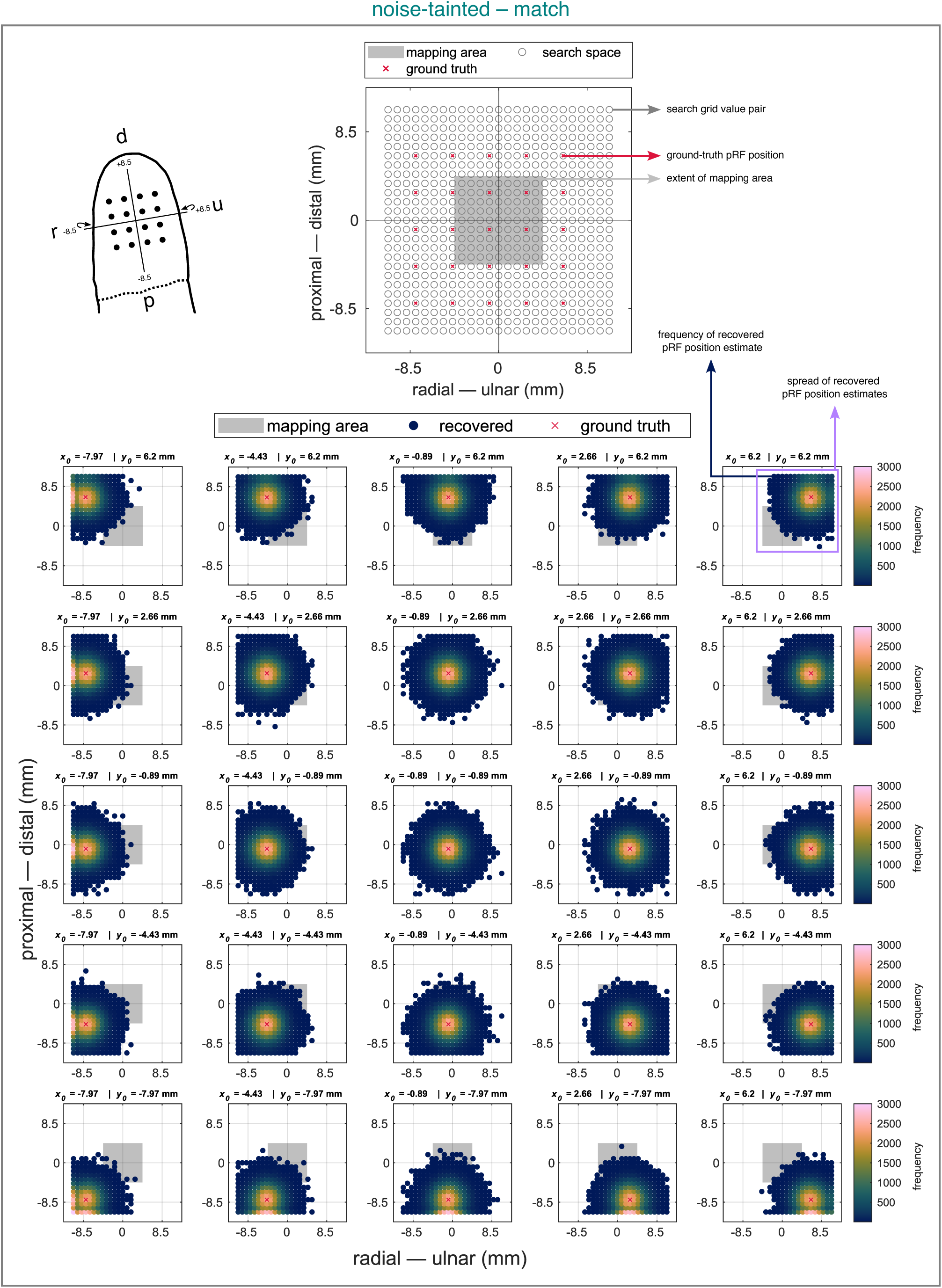
Recovered pRF position estimates for noise-tainted time series simulated and fit using the 2dg-fix model with ground-truth pRFs matching the search grid value pairs. Upper panel: Ground-truth pRF positions relative to search grid value pairs. Lower panels: Ground-truth pRF positions relative to recovered pRF position estimates. For each ground-truth pRF, 100000 repeats were simulated and the number of times a certain grid search value pair was recovered was counted. Note that although we simulated an odd and and even dataset (see text for details), for reasons of brevity, we only present the results of the even dataset here. Headers in each panel list ground-truth pRF parameters. Wrap-around arrows indicate an *x*-axis that is slightly wrapped around the fingertip. 2dg-fix = 2D Gaussian pRF model with parameter for pRF size fixed and constrained grid search. pRF = population receptive field. d = distal. p = proximal. r = radial. u = ulnar.

**Figure S4.**
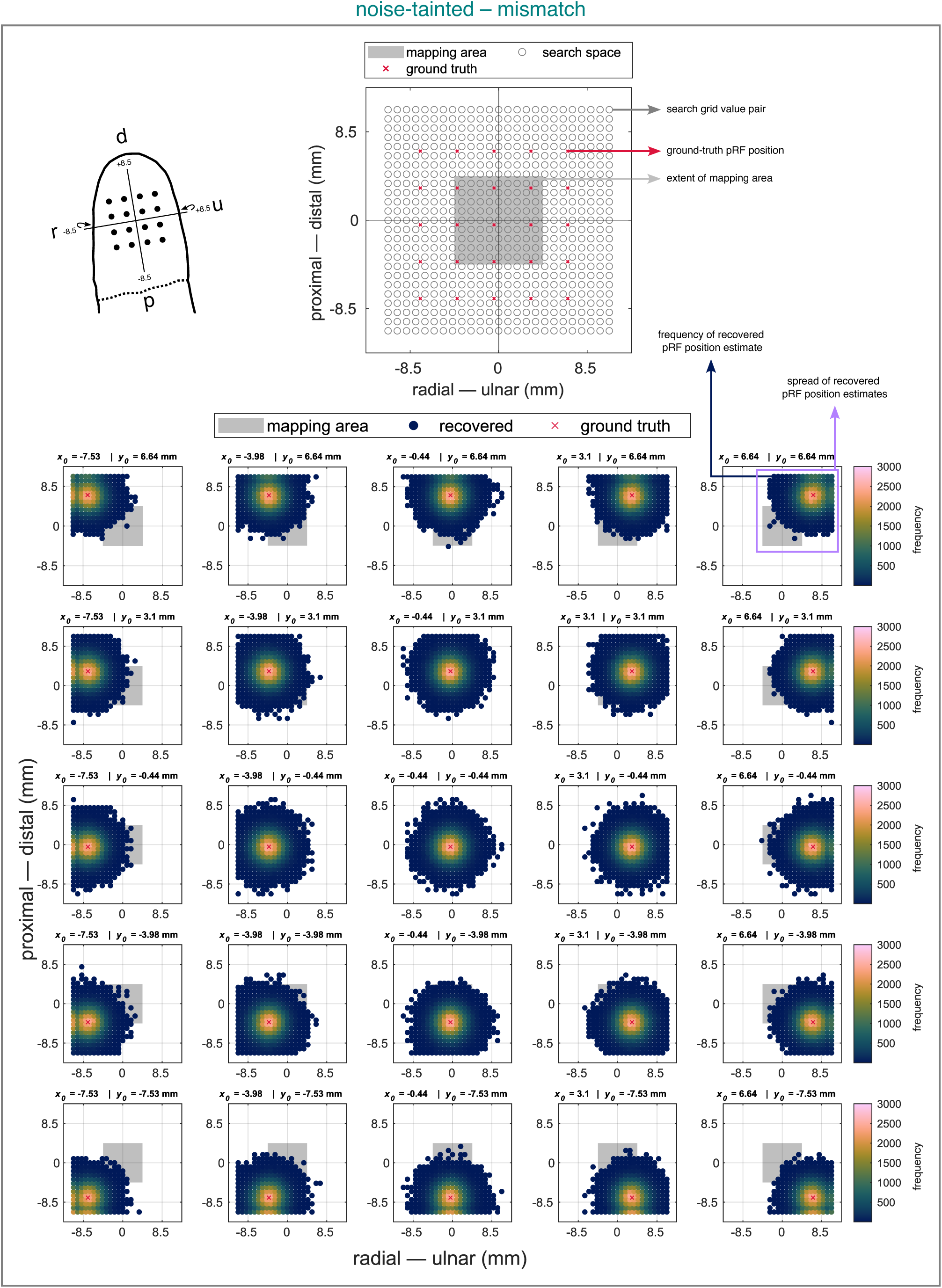
Recovered pRF positions estimates for noise-tainted time series simulated and fit using the 2dg-fix model with ground-truth pRFs mismatching the search grid value pairs. Upper panel: Ground-truth pRF positions relative to search grid value pairs. Lower panels: Ground-truth pRF positions relative to recovered pRF position estimates. For each ground-truth pRF, 100000 repeats were simulated and the number of times a certain grid search value pair was recovered was counted. Note that although we simulated an odd and and even dataset (see text for details), for reasons of brevity, we only present the results of the even dataset here. Headers in each panel list ground-truth pRF parameters. Wrap-around arrows indicate an *x*-axis that is slightly wrapped around the fingertip. 2dg-fix = 2D Gaussian pRF model with parameter for pRF size fixed and constrained grid search. pRF = population receptive field. d = distal. p = proximal. r = radial. u = ulnar.

**Figure S5.**
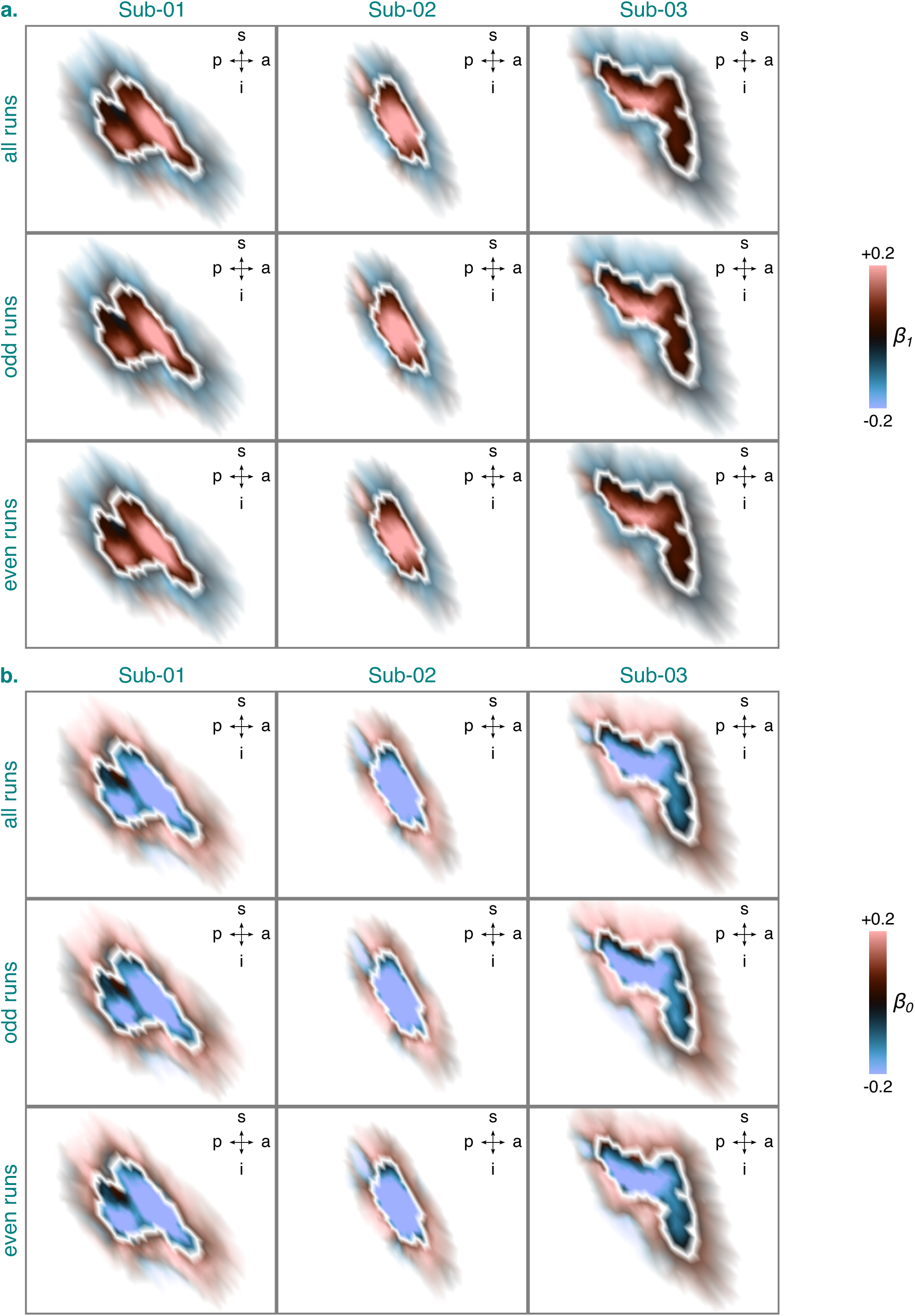
PRF amplitude (*β*_1_ in a.) and pRF baseline (*β*_0_ in b.) estimates for the identified BA3b cluster and the 2dg-fix model per participant and dataset. The *β*_0_ and *β*_1_ estimates are based on data from session 2-4. They were projected onto an inflated cortical surface model where dark gray regions represent sulci and light gray regions gyri. White lines denote the extent of the identified BA3b cluster (see Figure 4). A zoomed-in view is shown here showing the *β*_0_ and *β*_1_ estimates inside the identified BA3b cluster (opaque) and directly outside of it (semi-transparent). Any *β*_0_ or *β*_1_ estimates surpassing a value of ±0.2 were set to this value. BA3b = Brodmann area 3b. s = superior. p = posterior. i = inferior. a = anterior. Sub-01, Sub-02, Sub-03 = subject 01, 02, and 03. 2dg-fix = 2D Gaussian pRF model with parameter for pRF size fixed and constrained grid search. pRF = population receptive field.

**Figure S6.**
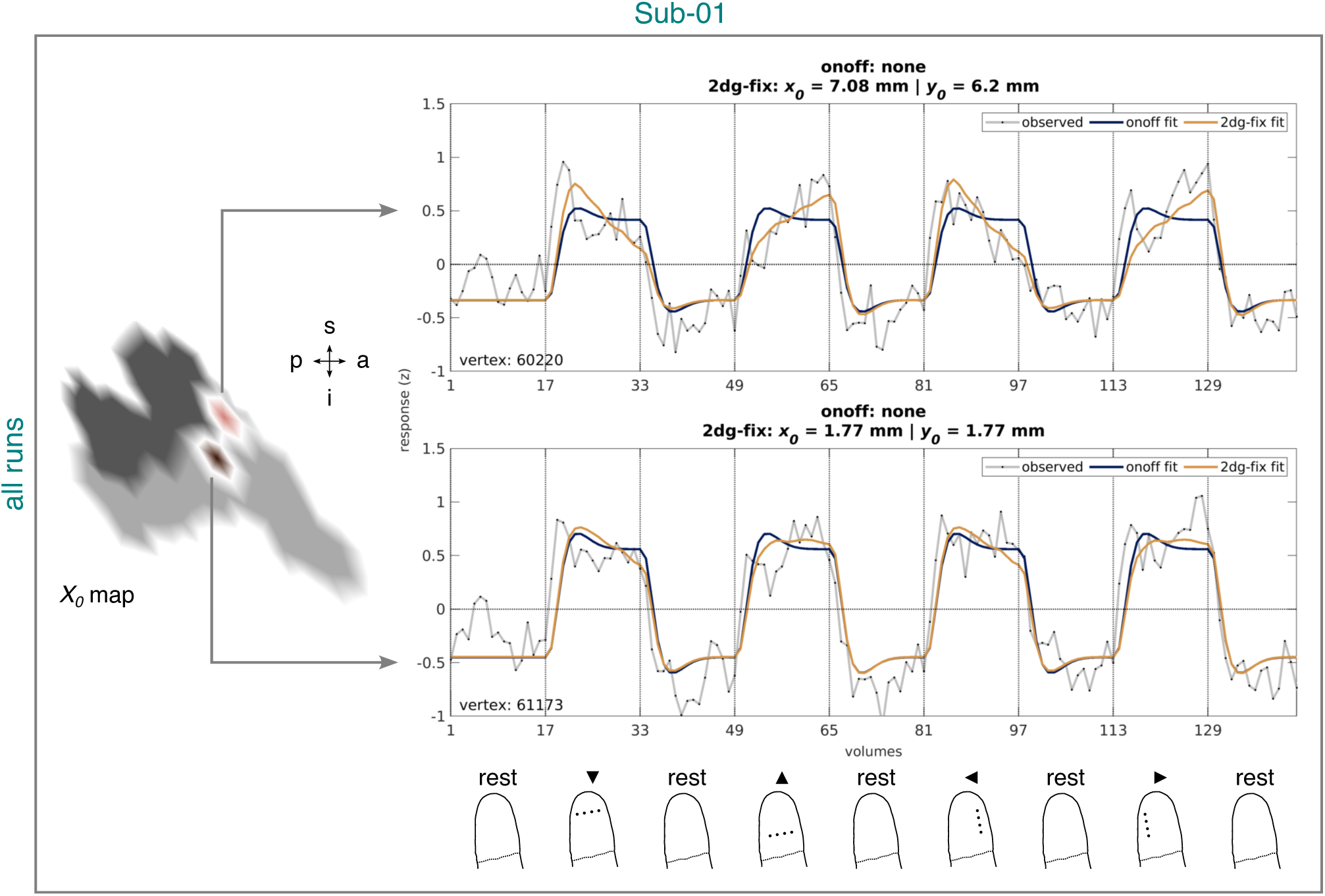
Observed and fitted time series corresponding to the onoff and 2dg-fix model from the identified BA3b cluster for Sub-01 and all runs. The time series are based on data from session 2-4. Left panel: Example vertices from the quantified *x*_0_ map projected onto an inflated cortical surface model where dark gray regions represent sulci and light gray regions gyri (see Figure 7, a.). White lines circumscribe the example vertices. A zoomed-in view is shown here, limited to the extent of the identified BA3b cluster (see Figure 4). Right upper and middle panel: Time series corresponding to the example vertices. Headers in each panel list pRF position estimates (if available) for each model. Vertex identifier is shown on the bottom left of each panel. Vertical lines accompanied by tick labels denote the onset of stimulation and baseline (rest) intervals. Right lower panel: Status of the fingertip at the onset of stimulation and baseline (rest) intervals (for more details, see Figure 2). Sub-01 = Subject 01. 2dg-fix = 2D Gaussian pRF model with parameter for pRF size fixed and constrained grid search. onoff = pRF model without any spatial tuning that simply reflects the presence or absence of tactile stimulation. pRF = population receptive field. BA3b = Brodmann area 3b. s = superior. p = posterior. i = inferior. a = anterior.

**Figure S7.**
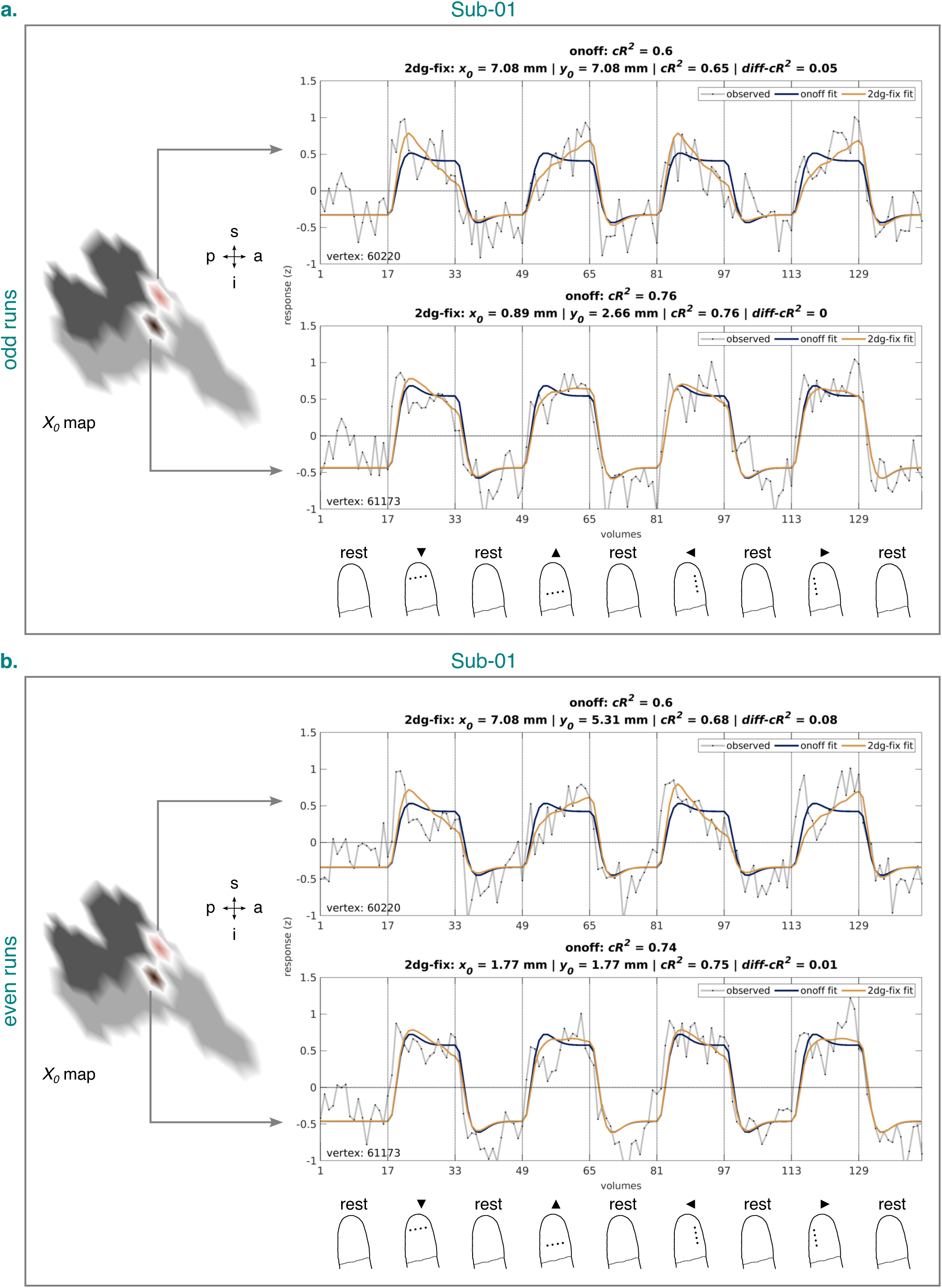
Observed and fitted time series corresponding to the onoff and 2dg-fix model from the identified BA3b cluster for Sub-01 and the odd (a.) and even (b.) runs. The time series are based on data from session 2-4. Left panel (in each subfigure): Example vertices from the quantified *x*_0_ map projected onto an inflated cortical surface model where dark gray regions represent sulci and light gray regions gyri (see Figure 7, a.). White lines circumscribe the example vertices. A zoomed-in view is shown here, limited to the extent of the identified BA3b cluster (see Figure 4). Right upper and middle panel (in each subfigure): Time series corresponding to the example vertices. Headers in each panel list pRF position estimates (if available) and cross-validated goodness-of-fit measures for each model. Vertex identifier is shown on the bottom left of each panel. Vertical lines accompanied by tick labels denote the onset of stimulation and baseline (rest) intervals. Right lower panel (in each subfigure): Status of the fingertip at the onset of stimulation and baseline (rest) intervals (for more details, see Figure 2). Sub-01 = Subject 01. 2dg-fix = 2D Gaussian pRF model with parameter for pRF size fixed and constrained grid search. onoff = pRF model without any spatial tuning that simply reflects the presence or absence of tactile stimulation. pRF = population receptive field. BA3b = Brodmann area 3b. s = superior. p = posterior. i = inferior. a = anterior.

**Figure S8.**
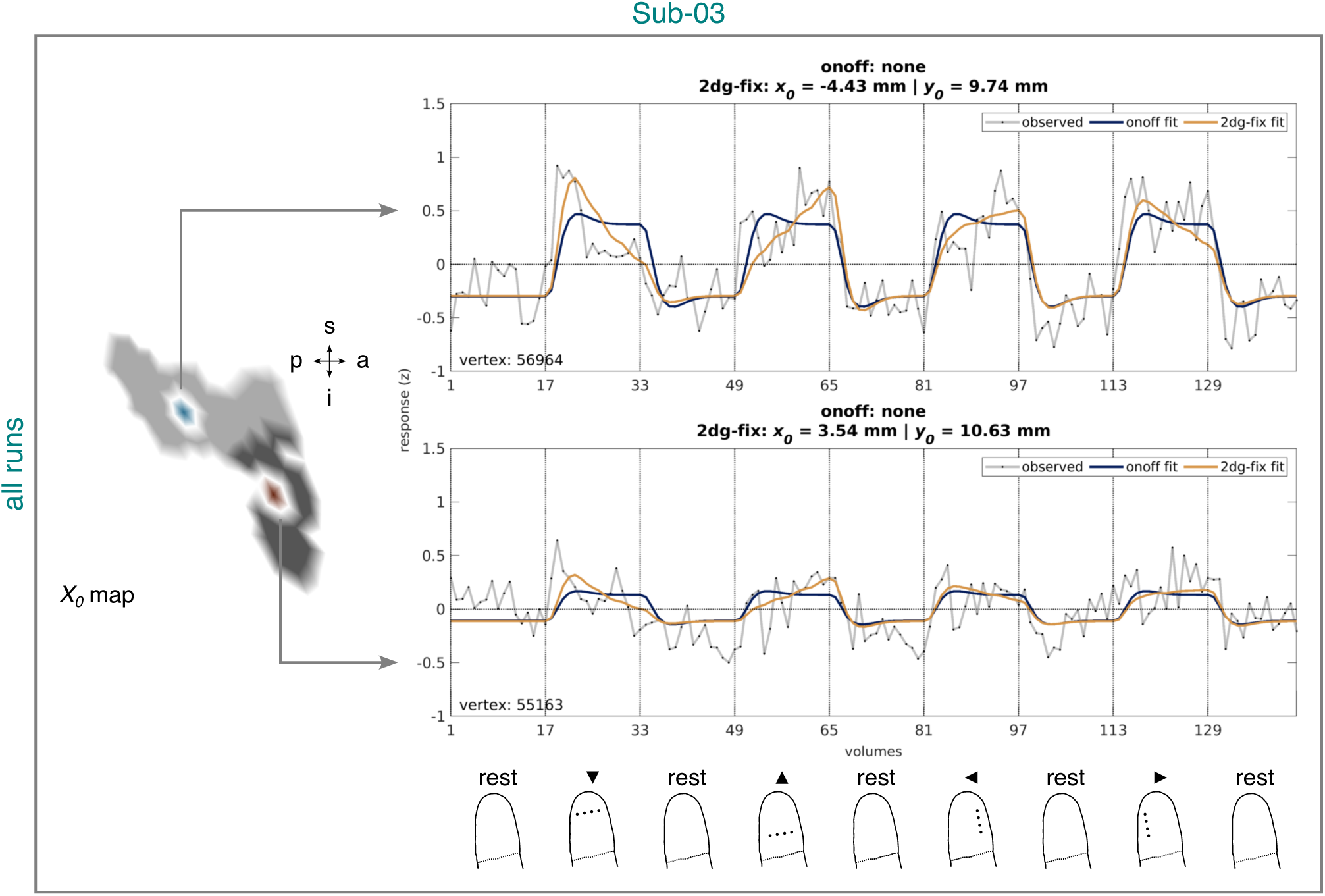
Observed and fitted time series corresponding to the onoff and 2dg-fix model from the identified BA3b cluster for Sub-03 and all runs. The time series are based on data from session 2-4. Left panel: Example vertices from the quantified *x*_0_ map projected onto an inflated cortical surface model where dark gray regions represent sulci and light gray regions gyri (see Figure 7, a.). White lines circumscribe the example vertices. A zoomed-in view is shown here, limited to the extent of the identified BA3b cluster (see Figure 4). Right upper and middle panel: Time series corresponding to the example vertices. Headers in each panel list pRF position estimates (if available) for each model. Vertex identifier is shown on the bottom left of each panel. Vertical lines accompanied by tick labels denote the onset of stimulation and baseline (rest) intervals. Right lower panel: Status of the fingertip at the onset of stimulation and baseline (rest) intervals (for more details, see Figure 2). Sub-03 = Subject 03. 2dg-fix = 2D Gaussian pRF model with parameter for pRF size fixed and constrained grid search. onoff = pRF model without any spatial tuning that simply reflects the presence or absence of tactile stimulation. pRF = population receptive field. BA3b = Brodmann area 3b. s = superior. p = posterior. i = inferior. a = anterior.

**Figure S9.**
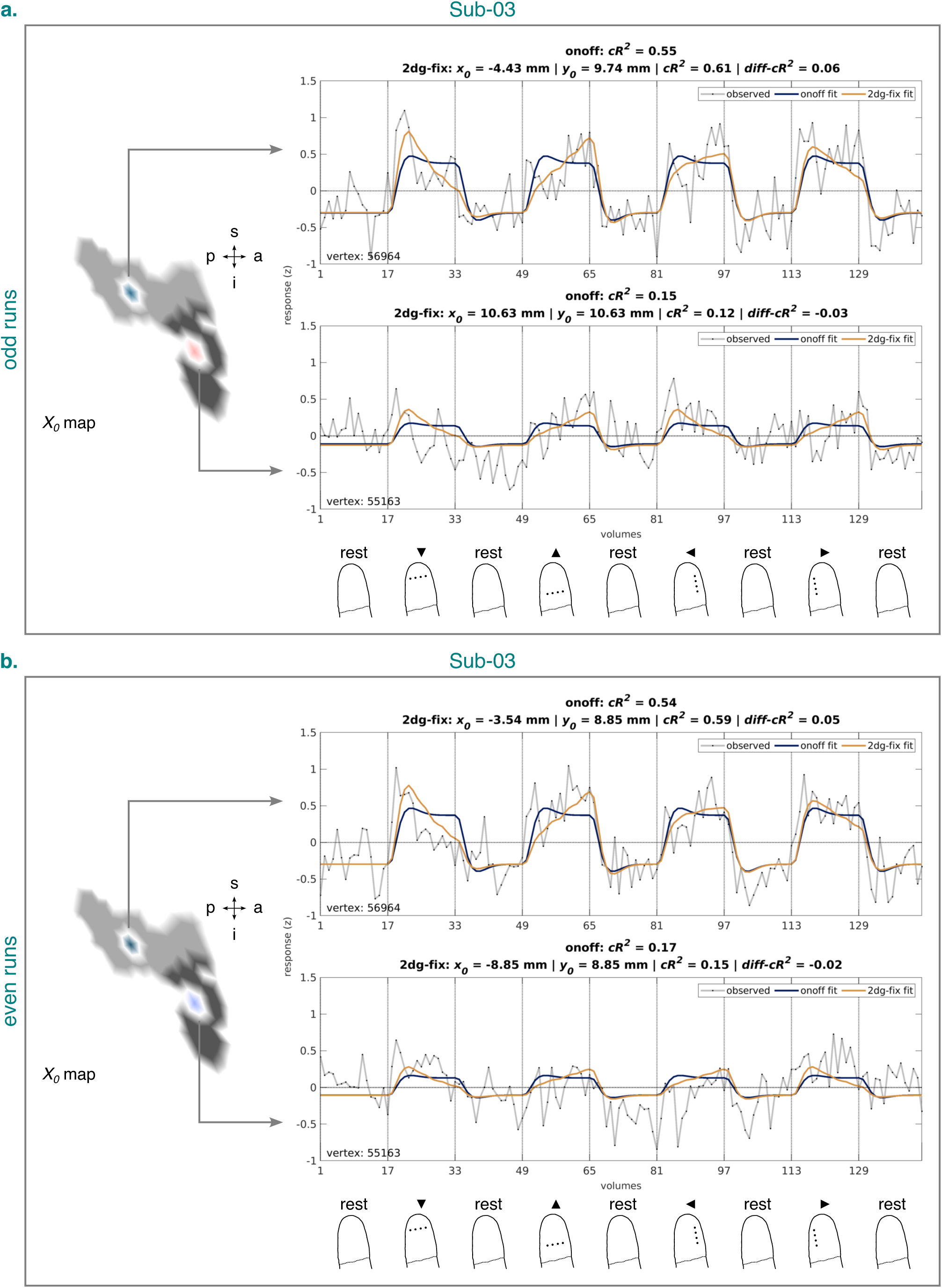
Observed and fitted time series corresponding to the onoff and 2dg-fix model from the identified BA3b cluster for Sub-03 and the odd (a.) and even (b) runs. The time series are based on data from session 2-4. Left panel (in each subfigure): Example vertices from the quantified *x*_0_ map projected onto an inflated cortical surface model where dark gray regions represent sulci and light gray regions gyri (see Figure 7, a.). White lines circumscribe the example vertices. A zoomed-in view is shown here, limited to the extent of the identified BA3b cluster (see Figure 4). Right upper and middle panel (in each subfigure): Time series corresponding to the example vertices. Headers in each panel list pRF position estimates (if available) and cross-validated goodness-of-fit measures for each model. Vertex identifier is shown on the bottom left of each panel. Vertical lines accompanied by tick labels denote the onset of stimulation and baseline (rest) intervals. Right lower panel (in each subfigure): Status of the fingertip at the onset of stimulation and baseline (rest) intervals (for more details, see Figure 2). Sub-03 = Subject 03. 2dg-fix = 2D Gaussian pRF model with parameter for pRF size fixed and constrained grid search. onoff = pRF model without any spatial tuning that simply reflects the presence or absence of tactile stimulation. pRF = population receptive field. BA3b = Brodmann area 3b. s = superior. p = posterior. i = inferior. a = anterior.

## Supplementary videos

*Video S1.* **BIDS-formatted anatomical image of Sub-02 in session 1.** The video runs through the whole volume and shows the image after preprocessing step 2 (see Figure S1). Sub-02 = subject 02. BIDS = Brain Imaging Data Structure.

*Video S2.* **Downsampled anatomical image of Sub-02 in session 1.** The video runs through the whole volume and shows the image after preprocessing step 3 (see Figure S1). Sub-02 = subject 02.

*Video S3.* **B1-bias-field-corrected anatomical image of Sub-02 in session 1.** The video runs through the whole volume and shows the image after preprocessing step 4 (see Figure S1). Sub-02 = subject 02.

*Video S4.* **Masked anatomical image of Sub-02 in session 1.** The video runs through the whole volume and shows the image after preprocessing step 5 (see Figure S1). Sub-02 = subject 02.

*Video S5.* **Initial masked anatomical template of Sub-02.** The video runs through the whole volume and shows the image after preprocessing step 6 (see Figure S1). Sub-02 = subject 02.

*Video S6.* **Unmasked anatomical template of Sub-02.** The video runs through the whole volume and shows the image after preprocessing step 7 (see Figure S1). Sub-02 = subject 02.

*Video S7.* **Final masked anatomical template of Sub-02.** The video runs through the whole volume and shows the image after preprocessing step 8 (see Figure S1). Sub-02 = subject 02.

*Video S8.* **BIDS-formatted functional image of Sub-02 in session 1 for run 1 and volume 1.** The video runs through the whole volume and shows the image after preprocessing step 2 (see Figure S1). Sub-02 = subject 02. BIDS = Brain Imaging Data Structure.

*Video S9.* **Average functional image of Sub-02 in session 1 for run 1.** The video runs through the whole volume and shows the image after preprocessing step 11 (see Figure S1). Sub-02 = subject 02.

*Video S10.* **Masked average functional image of Sub-02 in session 1 for run 1.** The video runs through the whole volume and shows the image after preprocessing step 13 (see Figure S1). Sub-02 = subject 02.

*Video S11.* **Masked functional template of Sub-02.** The video runs through the whole volume and shows the image after preprocessing step 14 (see Figure S1). Sub-02 = subject 02.

*Video S12.* **Coregistration: anatomical to functional for Sub-02.** The video shows the final masked anatomical template of Sub-02 coregistered to the masked functional template of Sub-02 for a single slice, and thus images after preprocessing step 15 and 14, respectively (see Figure S1). It loops 10 times through these images. Sub-02 = subject 02.

*Video S13.* **Coarse coregistration: functional to anatomical for Sub-02.** The video shows the BIDS-formatted functional image of Sub-02 in session 1 for run 1 and volume 1 coarsely coregistered to the final masked anatomical template of Sub-02 for a single slice, and thus images after preprocessing step 16 and 8, respectively (see Figure S1). It loops 10 times through these images. Sub-02 = subject 02. BIDS = Brain Imaging Data Structure.

*Video S14.* **Fine coregistration: functional to anatomical for Sub-02.** The video shows the coarsely-coregistered functional image of Sub-02 in session 1 for run 1 and volume 1 finely coregistered to the unmasked anatomical template (in this case FreeSurfer’s orig.mgz image) for a single slice, and thus images after preprocessing step 17 and 9, respectively (see Figure S1). It loops 10 times through these images. The blue line denotes the reconstructed gray-white matter boundary. Sub-02 = subject 02.

## Supplementary tables

**Table S1.**
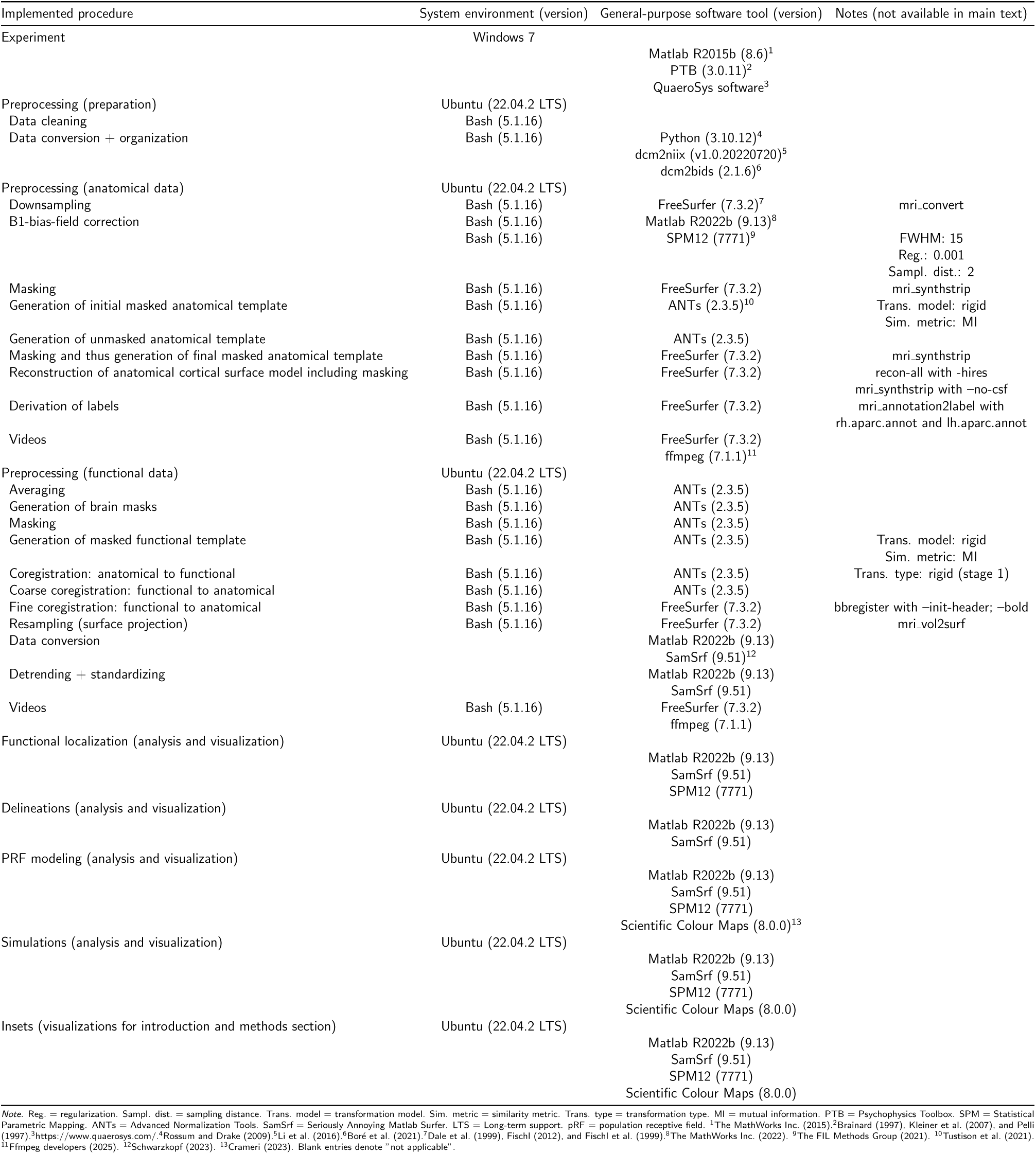
System Environment, General-Purpose Software Tool, and Notes as a Function of Implemented Procedure.

**Table S2.** Median Cross-validated Goodness-of-fit (*cR*^2^) as a Function of Participants/Simulation Type, Fitted pRF Model, and Cross-Validation Fold.

| Participants/simulation type | 2dg-fix |  | onoff |  |
| --- | --- | --- | --- | --- |
|  | odd on even | even on odd | odd on even | even on odd |
| Sub-01 | 0.39 | 0.35 | 0.36 | 0.32 |
| Sub-02 | 0.54 | 0.60 | 0.50 | 0.55 |
| Sub-03 | 0.20 | 0.20 | 0.19 | 0.21 |
| Sim<sub>2dg-fix-accuracy-match</sub> | 0.55 | 0.55 |  |  |
| Sim<sub>2dg-fix-accuracy-mismatch</sub> | 0.55 | 0.55 |  |  |
| Sim<sub>onoff-cross-validated-goodness-of-fit</sub> | 0.54 | 0.54 | 0.54 | 0.54 |

*Note.* pRF = population receptive field. 2dg-fix = 2D Gaussian pRF model with parameter for pRF size fixed and constrained grid search. onoff = pRF model without any spatial tuning that simply reflects the presence or absence of tactile stimulation. odd(even) on even(odd) = odd(even) runs or simulated data validated on even(odd) runs or simulated data. Sim<sub>2dg-fix-accuracy-match</sub> = data were simulated using the 2dg-fix model for ground-truth pRFs matching the search grid value pairs to assess the computational validity (accuracy) of pRF estimates. Sim<sub>2dg-fix-accuracy-mismatch</sub> = just like Sim<sub>2dg-fix-accuracy-match</sub>, but for ground-truth pRFs mismatching the search grid value pairs. Sim<sub>onoff-cross-validated-goodness-of-fit</sub> = data were simulated using the onoff model to assess the meaningfulness of differences in cross-validated goodness-of-fit. Blank entries denote "not applicable". Sub-01, Sub-02, Sub-03 = subject 01, 02, and 03.

## Footnotes

1 E.g. via a Fourier transform or cross-correlation procedure.

2 This means the data involved in functional localization and delineation (see 2.7.1 Functional localization and 2.7.2 Delineations) were independent of those involved in pRF modeling, thus circumventing issues of circularity (e.g., Kriegeskorte et al., 2009; Stoll et al., 2022).

3 It has been shown that averaging is particularly beneficial for heavily noise-tainted fMRI data both in terms of goodness-of-fit and accuracy of pRF parameter estimates (Senden et al., 2014), such as 7T-fMRI data at submillimeter resolution.

4 To determine this range, we excluded vertices with a ***t***-value ≤ 0. Considering that manual delineation based on ***t***-statistic maps (see 2.7.1 Functional localization and 3.1 Vibrotactile stimulation produces fingertip clusters in BA3b) is rarely perfectly precise, it can occasionally happen that vertices with ***t***-values ≤ 0 are included in a delineated cluster.

5 Importantly, however, the receptive fields measured in these studies were minimum receptive fields, defined as the area on the skin where light tactile stimulation evoked a maximal response.

6 To determine this range, we excluded vertices with a ***t***-value ≤ 0 and vertices for which at least one of the pRF models (2dg-fix or onoff) exhibited a negative cross-validated goodness-of-fit. The latter was done separately for each cross-validation fold. Considering that manual delineation based on ***t***-statistic maps (see 2.7.1 Functional localization and 3.1 Vibrotactile stimulation produces fingertip clusters in BA3b) is rarely perfectly precise, it can occasionally happen that vertices with ***t***-values ≤ 0 are included in a delineated cluster. Similarly, unlike traditional quantifications of goodness-of-fit using a single dataset, the cross-validated quantification adopted here can occasionally result in negative cross-validated goodness-of-fit values. Although these negative values are expected, they cannot be readily interpreted in the scope of the present modeling framework and were thus excluded.

